# Genome-wide characterization of satellite DNA arrays in a complex plant genome using nanopore reads

**DOI:** 10.1101/677575

**Authors:** Tihana Vondrak, Laura Ávila Robledillo, Petr Novák, Andrea Koblížková, Pavel Neumann, Jiří Macas

## Abstract

**Background:** Amplification of monomer sequences into long contiguous arrays is the main feature distinguishing satellite DNA from other tandem repeats, yet it is also the main obstacle in its investigation because these arrays are in principle difficult to assemble. Here we explore an alternative, assembly-free approach that utilizes ultra-long Oxford Nanopore reads to infer the length distribution of satellite repeat arrays, their association with other repeats and the prevailing sequence periodicities.

**Results:** We have developed a computational workflow for similarity-based detection and downstream analysis of satellite repeats in individual nanopore reads that led to genome-wide characterization of their properties. Using the satellite DNA-rich legume plant *Lathyrus sativus* as a model, we demonstrated this approach by analyzing eleven major satellite repeats using a set of nanopore reads ranging from 30 to over 200 kb in length and representing 0.73x genome coverage. We found surprising differences between the analyzed repeats because only two of them were predominantly organized in long arrays typical for satellite DNA. The remaining nine satellites were found to be derived from short tandem arrays located within LTR-retrotransposons that occasionally expanded in length. While the corresponding LTR-retrotransposons were dispersed across the genome, this array expansion occurred mainly in the primary constrictions of the *L. sativus* chromosomes, which suggests that these genome regions are favorable for satellite DNA accumulation.

**Conclusions:** The presented approach proved to be efficient in revealing differences in long-range organization of satellite repeats that can be used to investigate their origin and evolution in the genome.

## Background

Satellite DNA (satDNA) is a class of highly repeated genomic sequences characterized by its occurrence in long arrays of almost identical, tandemly arranged units called monomers. It is ubiquitous in animal and plant genomes, where it can make up to 36% or 18 Gbp/1C of nuclear DNA (Ambrožová *et al*., 2010). The monomer sequences are typically hundreds of nucleotides long, although they can be as short as simple sequence repeats (< 10 bp) (Heckmann *et al*., 2013) or reach over 5 kb (Gong *et al*., 2012). Thus, satDNA is best distinguished from other tandem repeats like micro- or minisatellites by forming much longer arrays (tens of kilobases up to megabases) that often constitute blocks of chromatin with specific structural and epigenetic properties (Garrido-Ramos, 2017). This genomic organization and skewed base composition have played a crucial role in satDNA discovery in the form of additional (satellite) bands observed in density gradient centrifugation analyses of genomic DNA (Kit, 1961). Thanks to a number of studies in diverse groups of organisms, the initial view of satellite DNA as genomic ‘junk’ has gradually shifted to an appreciation of its roles in chromosome organization, replication and segregation, gene expression, disease phenotypes and reproductive isolation between species (reviewed in Plohl et al., 2014; Garrido-Ramos, 2015, 2017; Hartley et al., 2019). Despite this progress, there are still serious limitations in our understanding of the biology of satDNA, especially with respect to the molecular mechanisms underlying its evolution and turnover in the genome.

Although the presence of satDNA is a general feature of eukaryotic genomes, its sequence composition is highly variable. Most satellite repeat families are specific to a single genus or even a species (Macas *et al*., 2002), which makes satDNA the most dynamic component of the genome. A theoretical framework for understanding satDNA evolution was laid using computer simulations (reviewed in Elder and Turner 1995). For example, the computer models demonstrated the emergence of tandem repeats from random non-repetitive sequences by a joint action of unequal recombination and mutation (Smith, 1976), predicted satDNA accumulation in genome regions with suppressed meiotic recombination (Stephan, 1986) and evaluated possible impacts of natural selection (Stephan & Cho, 1994). It was also revealed that recombination-based processes alone cannot account for the persistence of satDNA in the genome, which implied that additional amplification mechanisms need to be involved (Walsh, 1987). These models are of great value because, in addition to predicting conditions that can lead to satDNA origin, they provide testable predictions regarding tandem repeat homogenization patterns, the emergence of higher-order repeats (HORs) and the gradual elimination of satDNA from the genome. However, their utilization and further development have been hampered by the lack of genome sequencing data revealing the long-range organization and sequence variation within satDNA arrays that were needed to test their predictions.

A parallel line of research has focused on elucidating satDNA evolution using molecular and cytogenetic methods. These studies confirmed that satellite repeats can be generated by tandem amplification of various genomic sequences, for example, parts of dispersed repeats within potato centromeres (Gong *et al*., 2012) or a single-copy intronic sequence in primates (Valeri *et al*., 2018). An additional putative mechanism of satellite repeat origin was revealed in DNA replication studies, which showed that repair of static replication forks leads to the generation of tandem repeat arrays (Kuzminov, 2016). SatDNA can also originate by expansion of existing short tandem repeat arrays present within rDNA spacers (Macas *et al*., 2003) and in hypervariable regions of LTR-retrotransposons (Macas *et al*., 2009). Moreover, there may be additional links between the structure or transpositional activity of mobile elements and satDNA evolution (Meštrović *et al*., 2015). Once amplified, satellite repeats usually undergo a fast sequence homogenization within each family, resulting in high similarities of monomers within and between different arrays. This process is termed concerted evolution (Elder & Turner, 1995) and is supposed to employ various molecular mechanisms, such as gene conversion, segmental duplication and rolling-circle amplification of extrachromosomal circular DNA. However, little evidence has been gathered thus far to evaluate real importance of these mechanisms for satDNA evolution. Since each of these mechanisms leave specific molecular footprints, this question can be tackled by searching for these patterns within satellite sequences. However, obtaining such sequence data from a wide range of species has long been a limiting factor in satDNA investigation.

The introduction of next generation sequencing (NGS) technologies (Metzker, 2009) marked a new era in genome research, including the characterization of repetitive DNA (Weiss-Schneeweiss *et al*., 2015). Although the adoption of NGS resulted in a boom of genome assemblies, the genomes assembled using short-read technologies like Illumina are of limited use for satDNA investigation because they mostly lack satellite arrays (Peona *et al*., 2018). On the other hand, the short-read data are successfully utilized by bioinformatic pipelines specifically tailored to the identification of satellite repeats employing assembly-free algorithms (Novák *et al*., 2010, 2017; Ruiz-Ruano *et al*., 2016). Although these approaches proved to be efficient in satDNA identification and revealed a surprising diversity of satellite repeat families in some plant and animal species (Macas *et al*., 2015; Ruiz-Ruano *et al*., 2016; Ávila Robledillo *et al*., 2018), they, in principle, could not provide much insight into their large-scale arrangement in the genome. In this respect, the real breakthrough was recently made by the so-called long-read sequencing technologies that include the Pacific Biosciences and Oxford Nanopore platforms. Especially the latter has, due to its principle of reading the sequence directly from a native DNA strand during its passage through a molecular pore, a great potential to generate “ultra-long” reads reaching up to one megabase (van Dijk *et al*., 2018). Different strategies utilizing such long reads for satDNA investigation can be envisioned. First, they can be combined with other genome sequencing and mapping data to generate hybrid assemblies in which satellite arrays are faithfully represented and then analyzed. This approach has already been successfully used for assembling satellite-rich centromere of the human chromosome Y (Jain *et al*., 2018) and for analyzing homogenization patterns of satellites in *Drosphila melanogaster* (Weissensteiner *et al*., 2017). Alternatively, it should be possible to infer various features of satellite repeats by analyzing repeat arrays or their parts present in individual nanopore reads. Since only a few attempts have been made to adopt this strategy (Cechova & Harris, 2018) it has yet to be fully explored, which is the subject of the present study.

In this work, we aimed to characterize the basic properties of satellite repeat arrays in a genome-wide manner by employing bioinformatic analyses of long nanopore reads. As the model for this study, we selected the grass pea (*Lathyrus sativus* L.), a legume plant with a relatively large genome (6.52 Gbp/C) and a small number of chromosomes (2n =14) which are amenable to cytogenetic experiments. The chromosomes have extended primary constrictions with multiple domains of centromeric chromatin (meta-polycentric chromosomes) (Neumann *et al*., 2015, 2016) and well-distinguishable heterochromatin bands indicative of the presence of satellite DNA. Indeed, repetitive DNA characterization from low-pass genome sequencing data revealed that the *L. sativus* genome is exceptionally rich in tandem repeats that include 23 putative satDNA families, which combined represent 10.7% of the genome (Macas *et al*., 2015). Focusing on the fraction of the most abundant repeats, we developed a workflow for their detection in nanopore reads and subsequent evaluation of the size distributions of their arrays, their sequence homogenization patterns and their interspersion with other repetitive sequences. This work revealed surprising differences of the array properties between the analyzed repeats, which allowed their classification into two groups that differed in origin and amplification patterns in the genome.

## Data Description

For the present study, we chose a set of sixteen putative satellites with estimated genome proportions exceeding a threshold of 0.1% and reaching up to 2.6% of the *L. sativus* genome (Table 1). These sequences were selected as the most abundant from a broader set of 23 tandem repeats that were previously identified in *L. sativus* using graph-based clustering of Illumina reads (Macas *et al*., 2015). The clusters selected from this study were further analyzed using a TAREAN pipeline (Novák *et al*., 2017), which confirmed their annotation as satellite repeats and reconstructed consensus sequences of their monomers (Supplementary file 1). The monomers were 32 bp to 660 bp long and varied in their AT/GC content (46.3-76.6% AT). Mutual sequence similarities were detected between some of the monomers, which suggested that they represented variants (sub-families) of the same repeat family (Supplementary Fig. S1). These included three variants of the satellite families FabTR-51 and FabTR-53 and two variants of FabTR-52 (Table 1). Except for the FabTR-52 sequences, which were found to be up to 96% identical to the repeat pLsat described by (Ceccarelli *et al*., 2010), none of the satellites showed similarities to sequences in public sequence databases. We assembled a reference database of consensus sequences and additional sequence variants of all selected satellite repeats to be used for similarity-based detection of these sequences in the nanopore reads. The reference sequences were put into the same orientation to allow for evaluation of the orientation of the arrays in the nanopore reads.

**Table 1.** Characteristics of the investigated satellite repeats

| Satellite family | Monomer<br>[bp] | AT [%] | Genomic<br>abundance |  | FISH probe |
| --- | --- | --- | --- | --- | --- |
| *Subfamily* |  |  | [%] | [Mbp/1C] |  |
| **FabTR-2** | 49 | 71.4 | 1.700 | 110.8 | LASm3H1 |
| **FabTR-51** |  |  | 3.101 | 202.2 |  |
| *FabTR-51-LAS-A* | 80 | 46.3 | 2.500 | 163.0 | LASm1H1 |
| *FabTR-51-LAS-B* | 79 | 51.9 | 0.560 | 36.5 | LasTR6_H1 |
| *FabTR-51-LAS-C* | 118 | 50.0 | 0.041 | 2.7 |  |
| **FabTR-52** |  |  | 2.019 | 131.6 |  |
| *FabTR-52-LAS-A* | 55 | 47.3 | 2.000 | 130.4 | LASm2H1 |
| *FabTR-52-LAS-B* | 32 | 50.0 | 0.019 | 1.2 |  |
| **FabTR-53** |  |  | 2.600 | 169.5 | c1644 + c1645 |
| *FabTR-53-LAS-A* | 660 | 76.6 | n.d. |  |  |
| *FabTR-53-LAS-B* | 368 | 76.4 | n.d. |  |  |
| *FabTR-53-LAS-C* | 565 | 75.9 | n.d. |  |  |
| **FabTR-54** | 104 | 51.0 | 0.840 | 54.8 | LasTR5_H1 |
| **FabTR-55** | 78 | 55.1 | 0.480 | 31.3 | LasTR7_H1 |
| **FabTR-56** | 46 | 60.9 | 0.250 | 16.3 | LasTR8_H1 |
| **FabTR-57** | 61 | 65.6 | 0.130 | 8.5 | LasTR9_H1 |
| **FabTR-58** | 86 | 59.3 | 0.140 | 9.1 | LasTR10_H1 |
| **FabTR-59** | 131 | 49.6 | 0.110 | 7.2 | LasTR11_H1 |
| **FabTR-60** | 86 | 52.3 | 0.110 | 7.2 | LasTR12_H1 |

We conducted two sequencing runs on the Oxford Nanopore MinION device utilizing independent libraries prepared from partially fragmented genomic DNA using a 1D ligation sequencing kit (SQK-LSK109). The two runs resulted in similar size distributions of the reads (Supplementary Fig. S2, panel A) and combined produced a total of 8.96 Gbp of raw read data. Following quality filtering, the reads shorter than 30 kb were discarded because we aimed to analyze only a fraction of the longest reads. The remaining 78,563 reads ranging from 30 kb to 348 kb in length (N50 = 67 kb) provided a total of 4.78 Gbp of sequence data, which corresponded to 0.73x coverage of the *L. sativus* genome.

## Analyses

### Detection of the satellite arrays in nanopore reads revealed repeats with contrasting array length distributions

The strategy for analyzing the length distribution of the satellite repeat arrays in the genome using nanopore reads is schematically depicted in Fig. 1. The satellite arrays in the nanopore reads were identified by similarity searches against the reference database employing the LASTZ program (Harris, 2007). Using a set of nanopore reads with known repeat compositions, we first optimized the LASTZ parameters towards high sensitivity and specificity. Under these conditions, the satDNA arrays within nanopore reads typically produced a series of short overlapping similarity hits that were filtered and parsed with custom scripts to detect the contiguous repeat regions longer than 300 bp. Then, the positions and orientations of the detected repeats were recorded, while distinguishing whether they were complete or truncated by the read end. In the latter case, the recorded array length was actually an underestimation of the real size.

**Figure 1.**
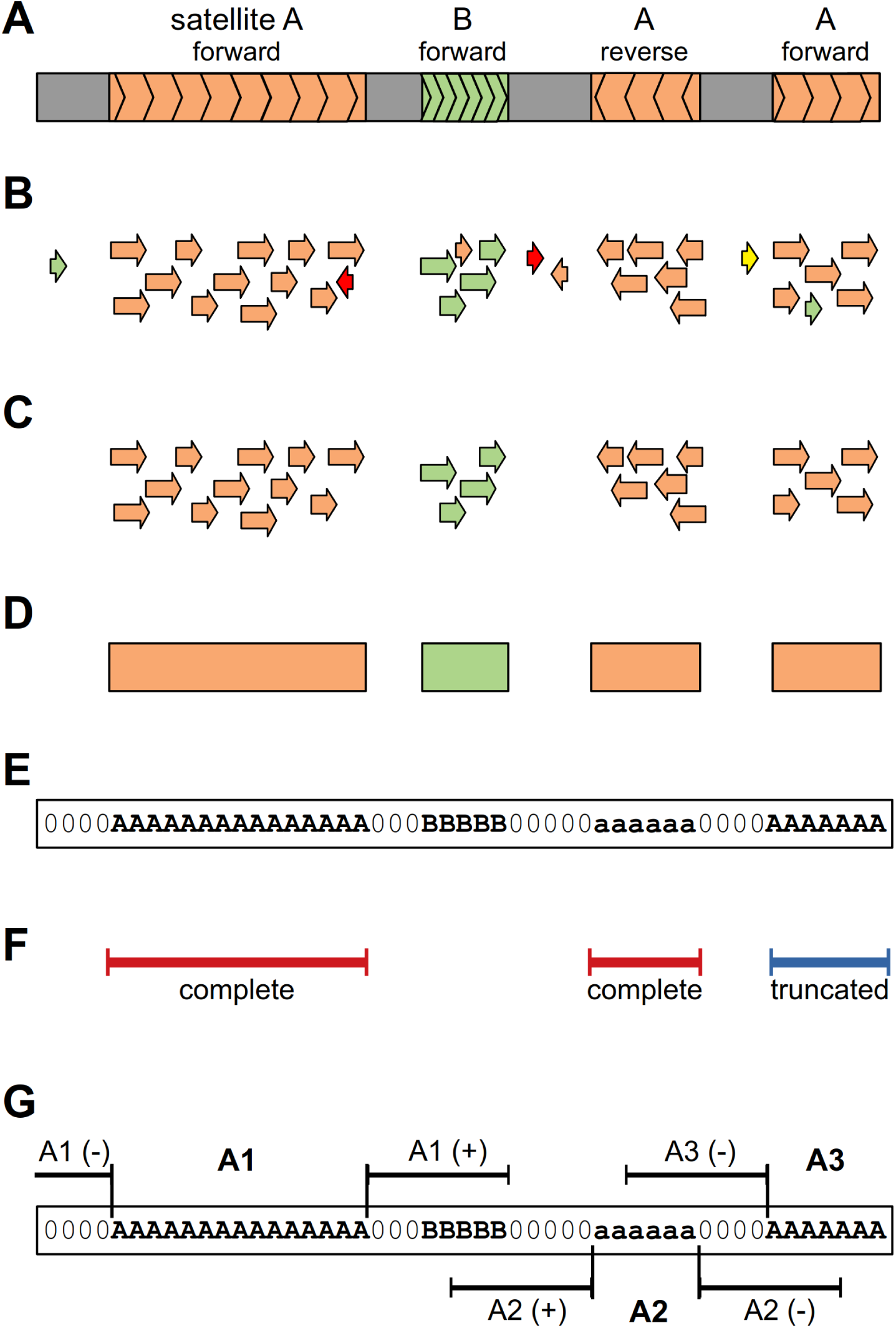
Schematic representation of the analysis strategy. (**A**) Nanopore read (gray bar) containing arrays of satellites A (orange) and B (green). The orientations of the arrays with respect to sequences in the reference database are indicated. (**B**) LASTZ search against the reference database results in similarity hits (displayed as arrows showing their orientation, with colors distinguishing satellite sequences) that are quality-filtered to remove non-specific hits (**C**). The filtered hits are used to identify the satellite arrays as regions of specified minimal length that are covered by overlapping hits to the same repeat (**D**). The positions of these regions are recorded in the form of coded reads where the sequences are replaced by satellite codes and array orientations are distinguished using uppercase and lowercase characters (**E**). The coded reads are then used for various downstream analyses. (**F**) Array lengths are extracted and analyzed regardless of orientation of the arrays but while distinguishing the complete and truncated arrays (here it is shown for satellite A). (**G**) Analysis of the sequences adjacent to the satellite arrays includes 10 kb regions upstream (-) and downstream (+) of the array. This analysis is performed with respect to the array orientation (compare the positions of upstream and downstream regions for arrays in forward (A1, A3) versus reverse orientation (A2)).

When the above analyses were applied to the whole set of nanopore reads, the detected array lengths were pooled for each satellite repeat, and their distributions were visualized as weighted histograms with a bin size of 5 kb, distinguishing complete and truncated satellite arrays (Fig. 2). This type of visualization accounts for the total lengths of the satellite sequences that occur in the genome as arrays of the lengths specified by the bins. Alternatively, the array size distributions were also plotted as histograms of their counts (Supplementary Fig. S3). As a control for the satellite repeats, we also analyzed the length distribution of 45S rDNA sequences, which typically form long arrays of tandemly repeated units (Copenhaver & Pikaard, 1996). Indeed, the plots revealed that most of the 45S rDNA repeats were detected as long arrays ranging up to >120 kb (Fig. 2). A similar pattern was expected for the satellite repeats; however, it was found for only two of them, FabTR-2 and FabTR-53. Both of these repeats were almost exclusively present as long arrays that extended beyond the lengths of most of the reads. To verify these results, we analyzed randomly selected reads using sequence self-similarity dot-plots, which confirmed that most of the arrays spanned entire reads or were truncated at only one of their ends (Supplementary Fig. S4 A,E). However, all nine remaining satellites generated very different array length distribution profiles that consisted of relatively large numbers of short (< 5 kb) arrays and comparatively fewer longer arrays (Fig. 2 and Supplementary Fig. S3). The proportions of these two size classes differed between the satellites, for example, while for FabTR-58, most of the arrays (98%) were short and only a few were expanded over 5 kb, FabTR-51 displayed a gradient of sizes from < 5 kb to 174 kb. To check whether these profiles could have partially been due to differences in the lengths of the reads containing these satellites, we also analyzed their size distributions. However, the read length distributions were similar between the different repeats, and there was no bias towards shorter read lengths (Supplementary Fig. S2, panel B). Thus, we concluded that nine of eleven analyzed satellites occurred in the *L. sativus* genome predominantly as short tandem arrays, and only a fraction of them expanded to form long arrays typical of satellite DNA. This conclusion was also confirmed by the dot-plot analyses of the individual reads, which revealed reads carrying short or intermediate-sized arrays and a few expanded ones (Supplementary Fig. S4 I-N).

**Figure 2.**
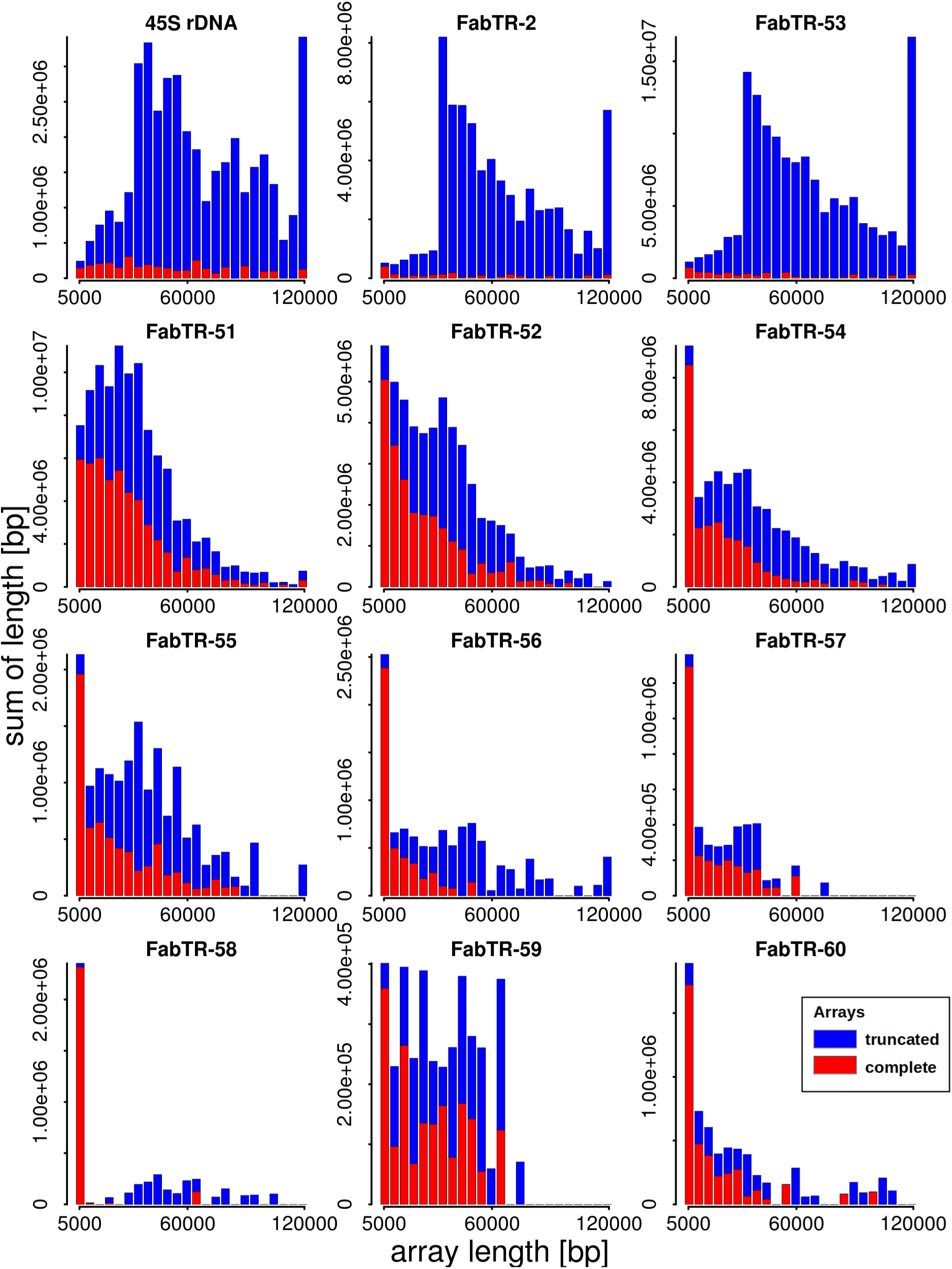
Length distributions of the satellite repeat arrays. The lengths of the arrays detected in the nanopore reads are displayed as weighted histograms with a bin size of 5 kb; the last bin includes all arrays longer than 120 kb. The arrays that were completely embedded within the reads (red bars) are distinguished from those that were truncated by their positions at the ends of the reads (blue bars). Due to the array truncation, the latter values are actually underestimations of the real lengths of the corresponding genomic arrays and should be considered as lower bounds of the respective array lengths.

### Analysis of genomic sequences adjacent to the satellite arrays identified a group of satellites that originated from LTR-retrotransposons

Next, we were interested in whether the investigated satellites were frequently associated in the genome with each other or with other types of repetitive DNA. Using a reference database for the different lineages of LTR-retrotransposons, DNA transposons, rDNA and telomeric repeats compiled from *L. sativus* repeated sequences identified in our previous study (Macas *et al*., 2015), we detected these repeats in the nanopore reads using LASTZ along with the analyzed satellites. Their occurrences were then analyzed within 10-kb regions directly adjacent to each satellite repeat array, and the frequencies at which they were associated with individual satDNA families were plotted with respect to the oriented repeat arrays (Fig. 3). When performed for the control 45S rDNA, this analysis revealed that they were mostly surrounded by arrays of the same sequences oriented in the same direction. This pattern emerged due to short interruptions of otherwise longer arrays. Similar results were found for FabTR-2 and FabTR-53 which also formed long arrays in the genome. Notably, the adjacent regions could be analyzed for only 33% and 35% of the FabTR-2 and FabTR-53 arrays, respectively, because these repeats mostly spanned entire reads. Substantially different profiles were obtained for the remaining nine satellites, revealing their frequent association with Ogre LTR-retrotransposons. No other repeats were detected at similar frequencies, except for unclassified LTR retrotransposons that probably represented less-conserved Ogre sequences. At a much smaller frequency (∼0.1), the FabTR-54 repeat was found to be adjacent to the FabTR-56 satellite arrays. Based on its position and size in relation to FabTR-56, the detected pattern corresponded to short FabTR-54 arrays attached to FabTR-56 in a direction-specific manner. Inspection of the individual reads confirmed that short arrays of these satellites occurred together in a part of the reads (Supplementary Fig. S4L). A peculiar pattern was revealed for FabTR-58 that consisted of a series of peaks that suggested interlacing FabTR-58 and Ogre sequences at fixed intervals (Fig. 3). This pattern was found to be due to occurrence of complex arrays consisting of multiple short arrays of FabTR-58 arranged in the same orientation and embedded into Ogre sequences (Supplementary Fig. S4Q). Upon closer inspection, this organization was found in numerous reads.

**Figure 3.**
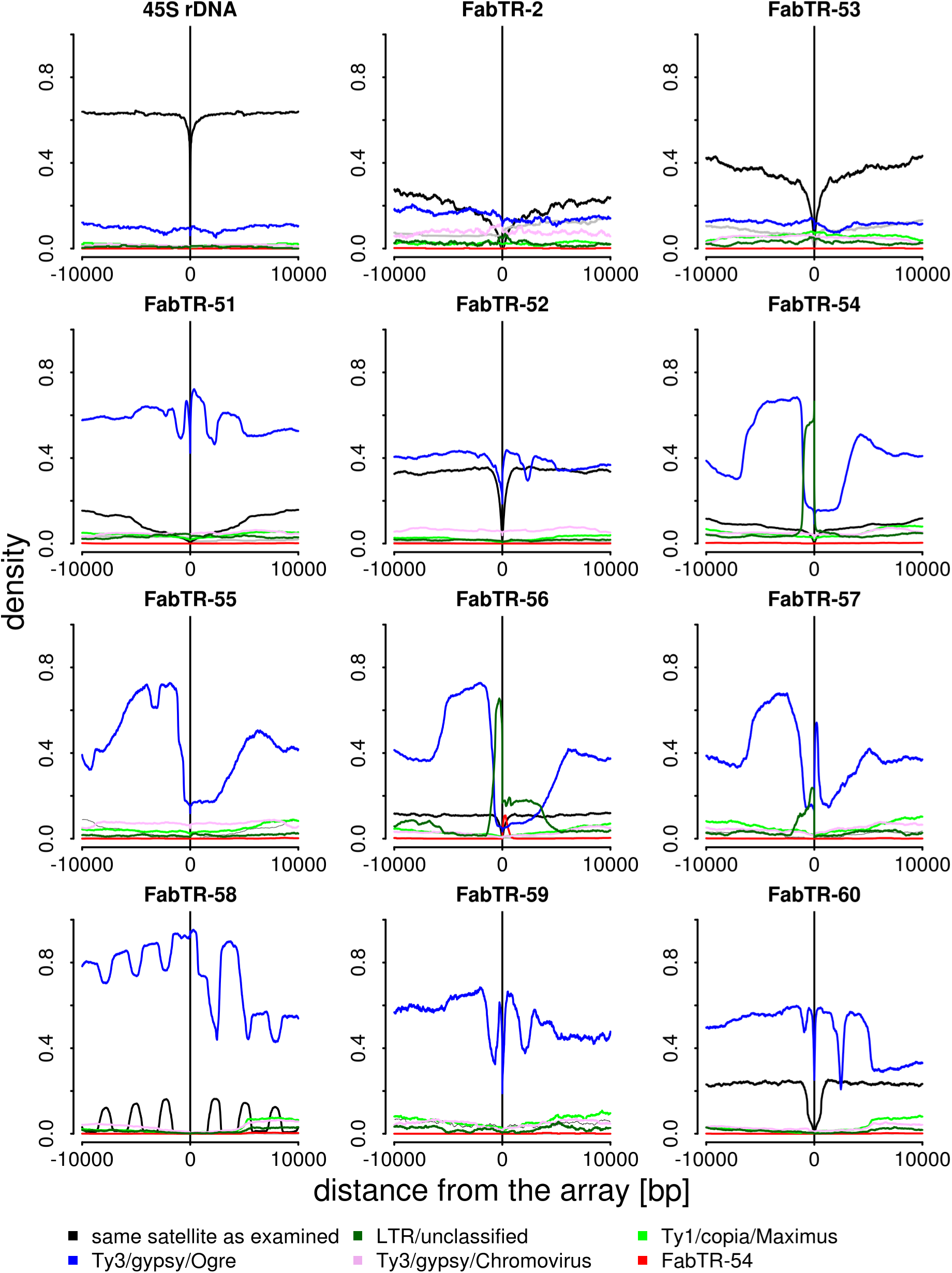
Sequence composition of the genomic regions adjacent to the satellite repeat arrays. The plots show the proportions of repetitive sequences identified within 10 kb regions upstream (positions −1 to −10,000) and downstream (1 to 10,000) of the arrays of individual satellites (the array positions are marked by vertical lines, and the plots are related to the forward-oriented arrays). Only the repeats detected in proportions exceeding 0.05 are plotted (colored lines). The black lines represent the same satellite as examined.

Ogre elements represent a distinct phylogenetic lineage of Ty3/gypsy LTR-retrotransposons (Neumann *et al*., 2019) that were amplified to high copy numbers in some plant species including *L. sativus*. Because they comprise 45% of the *L. sativus* genome (Macas *et al*., 2015), the frequent association of Ogres with short array satellites could simply be due to their random interspersion. However, we noticed from the structural analysis of the reads that these short arrays were often surrounded by two direct repeats, which is a feature typical of LTR-retrotransposons. This finding could mean that the arrays are actually embedded within the Ogre elements and were not only frequently adjacent to them by chance. To test this hypothesis, we performed an additional analysis of the array neighborhoods, but this time, we specifically detected parts of the Ogre sequences coding for the retroelement protein domains GAG, protease (PROT), reverse transcriptase (RT), RNase H (RH), archeal RNase H (aRH) and integrase (INT). If the association of Ogre sequences with the satellite arrays was random, these domains would be detected at various distances and orientations with respect to the arrays. In contrast, finding them in a fixed arrangement would confirm that the tandem arrays were in fact parts of the Ogre elements and occurred there in specific positions. As evident from Fig. 4A, that latter explanation was confirmed for all nine satellites. We found that their arrays occurred downstream of the Ogre *gag-pol* region including the LTR-retrotransposon protein coding domains in the expected order and orientation (see the element structure in Fig. 4B). In two cases (FabTR-54 and 57), some protein domains were not detected, and major peaks corresponded to the GAG domain which was relatively close to the tandem arrays. These patterns were explained by the frequent occurrence of these tandem arrays in non-autonomous elements lacking their *pol* regions due to large deletions. In approximately half of the satellites (*e.g.,* FabTR-51 and 52), we detected additional smaller peaks corresponding to the domains in both orientations located approximately 7-10 kb from the arrays. Further investigation revealed that these peaks represented Ogre elements that were inserted into the expanded arrays of corresponding satellites (Supplementary Fig. S4K). Consequently, they were detected only in satellites such as FabTR-51 and 52 in which the proportions of expanded arrays were relatively large and not FabTR-58 in which the expanded arrays were almost absent.

**Figure 4.**
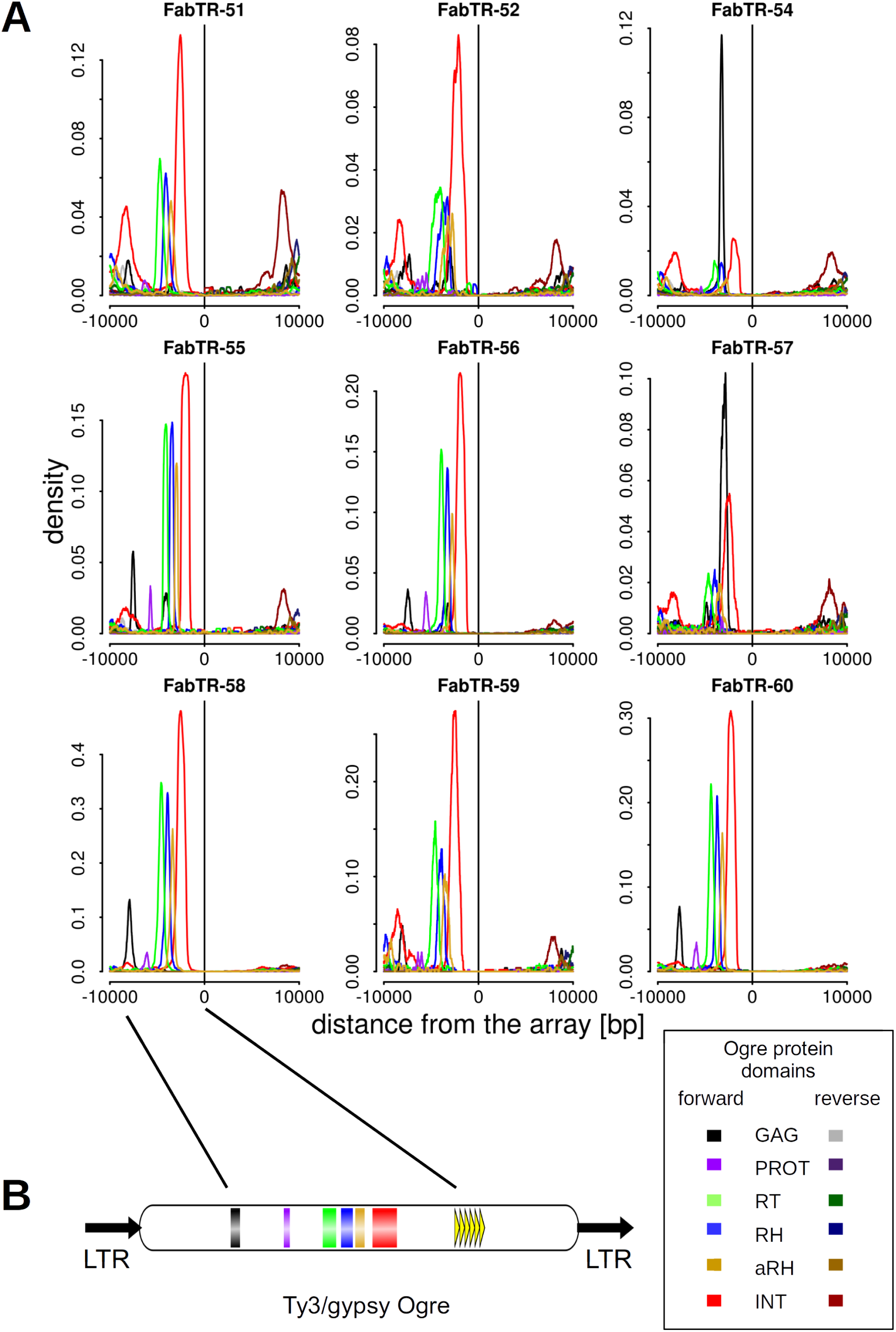
Detection of the Ogre sequences coding for the retrotransposon conserved protein domains in the genomic regions adjacent to the satellite repeat arrays. (**A**) The plots show the proportions of similarity hits from the individual domains and their orientation with respect to the forward-oriented satellite arrays. (**B**) A schematic representation of the Ogre element with the positions of the protein domains and short tandem repeats downstream of the coding region.

### Satellites with mostly expanded arrays show higher variation in their sequence periodicities

The identification of large numbers of satellite arrays in the nanopore reads provided sequence data for investigating the conservation of monomer lengths and the eventual occurrence of additional monomer length variants and HORs. To this purpose we designed a computational pipeline that extracted all satellite arrays longer than 30 kb and subjected them to a periodicity analysis using the fast Fourier transform algorithm (Venables & Ripley, 2002). The analysis revealed the prevailing monomer sizes and eventual additional periodicities in the tandem repeat arrays as periodicity spectra containing peaks at positions corresponding to the lengths of the tandemly repeated units. These periodicity spectra were averaged for all arrays of the same satellite (Fig. 5) or plotted separately for the individual arrays to explore the periodicity variations (Supplementary Fig. S5). As an alternative approach, we also visualized the array periodicities using nucleotide autocorrelation functions (Herzel *et al*., 1999; Macas *et al*., 2006). In selected cases, we verified the periodicity patterns within arrays using dot-plot analyses (Supplementary Fig. S4 B-D and F-H).

**Figure 5.**
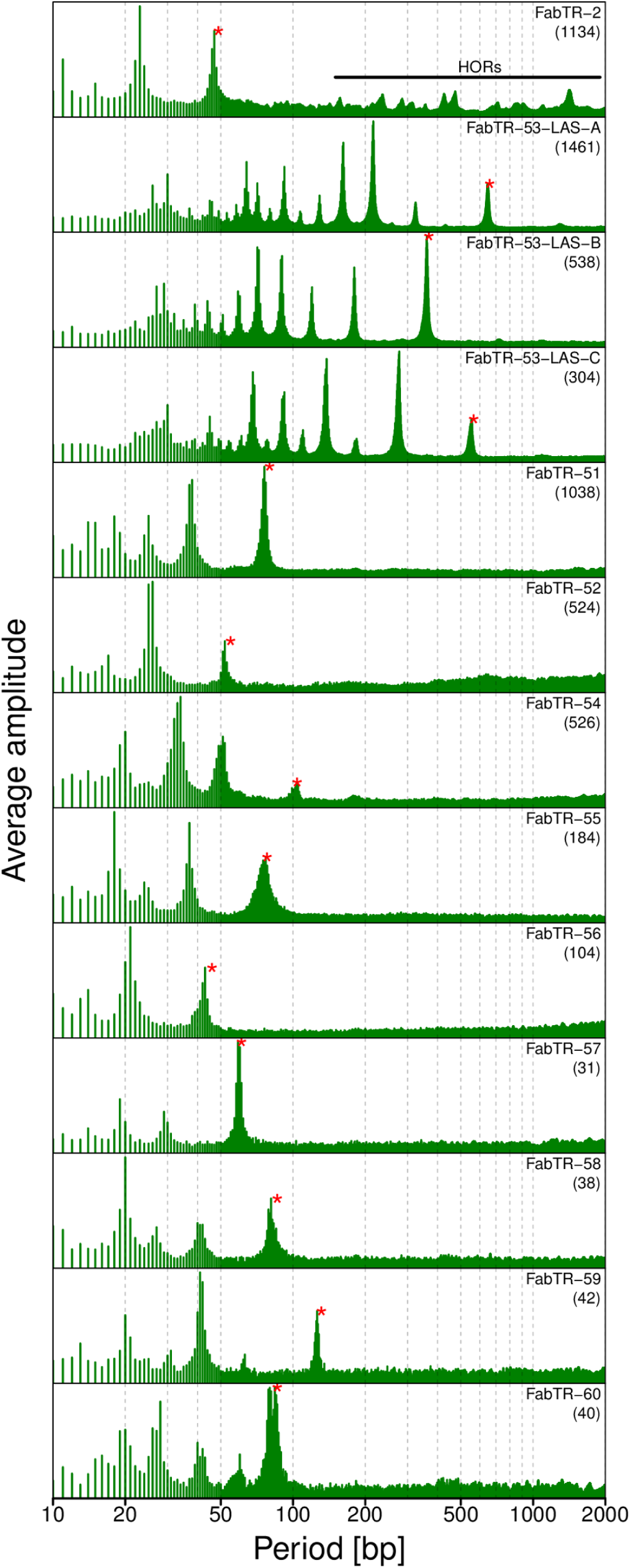
Periodicity spectra revealed by the fast Fourier transform analysis of the satellite repeat arrays. Each spectrum is an average of the spectra calculated for the individual arrays longer than 30kb of the same satellite family or subfamily. The numbers of arrays used for the calculations are in parentheses. The peaks corresponding to the monomer lengths listed in Table 1 are marked with red asterisks. The peaks in the FabTR-2 spectrum corresponding to higher-order repeats are indicated by the horizontal line.

As expected, the periodicity spectra of all satellites contained peaks corresponding to their monomer lengths (Fig. 5 and Table 1). In the nine Ogre-derived satellite repeats, the monomer periods were the longest detected and corresponded to the fundamental frequencies. There were only a few additional peaks detected with shorter periods that corresponded to higher harmonics (see Methods) or possibly reflected short subrepeats or underlying single-base periodicities. In contrast, FabTR-2 and FabTR-53 repeats, which occur in the genome as the expanded arrays, displayed more periodicity variations. Various HORs that probably originated from multimers of the 49 bp consensus were detected in the FabTR-2 arrays. Closer examination of the individual arrays revealed that the multiple peaks evident in the averaged periodicity spectrum (Fig. 5) originated as combinations of several simpler HOR patterns that differed between individual satellite arrays (Supplementary Fig. S5). In FabTR-53, the HORs were not detected, but a number shorter periodicities were revealed, which suggests that the current monomers of 660, 368 and 565 bp (subfamilies A, B and C, respectively) actually originated as higher-order repeats of shorter units of ∼190 bp (Fig. 5). An additional analysis using autocorrelation functions generally agreed with the fast Fourier transform approach and confirmed the high variabilities in FabTR-2 and FabTR-53 (Supplementary Fig. S5).

### Array expansion of the retrotransposon-derived satellites occurred preferentially in the pericentromeric regions of L. sativus chromosomes

To complement the analysis of satellite arrays with the information about their genomic distribution, we performed their detection on metaphase chromosomes using fluorescence *in situ* hybridization (FISH) (Fig. 6). Labeled oligonucleotides corresponding to the most conserved parts of the monomer sequences were used as hybridization probes in all cases except for FabTR-53 for which a mix of two cloned probes was used instead due to its relatively long monomers (Table 1 and Supplementary file 2). Although each satellite probe generated a different labeling pattern, most of them were located within the primary constrictions. The exception was FabTR-53, which produced strong hybridization signals that overlapped with most of the subtelomeric heterochromatin bands (Fig. 6A). The other distinct pattern was revealed for FabTR-2, which produced a series of dots along the periphery of the primary constrictions on all chromosomes (Fig. 6B). This pattern was identical to that obtained using an antibody to centromeric histone variant CenH3 (Neumann *et al*., 2015, 2016), which suggests that FabTR-2 is the centromeric satellite. The remaining nine probes corresponding to Ogre-derived satellites mostly produced bands at various parts of primary constrictions (Fig. 6C-F and Supplementary Fig. S6). For example, the bands of FabTR-54 occurred within or close to the primary constrictions of all chromosomes and produced a labeling pattern which, together with the chromosome morphology, allowed us distinguish all chromosome types within the *L. sativus* karyotype (Fig. 6C). A peculiar pattern was generated by the FabTR-51 subfamily A probe, which painted whole primary constrictions of one pair of chromosomes (chromosome 1, Fig. 6D); a similar pattern was produced by the FabTR-52 probe, but it labeled the entire primary constrictions of a different pair (chromosome 7, Fig. 6E).

**Figure 6.**
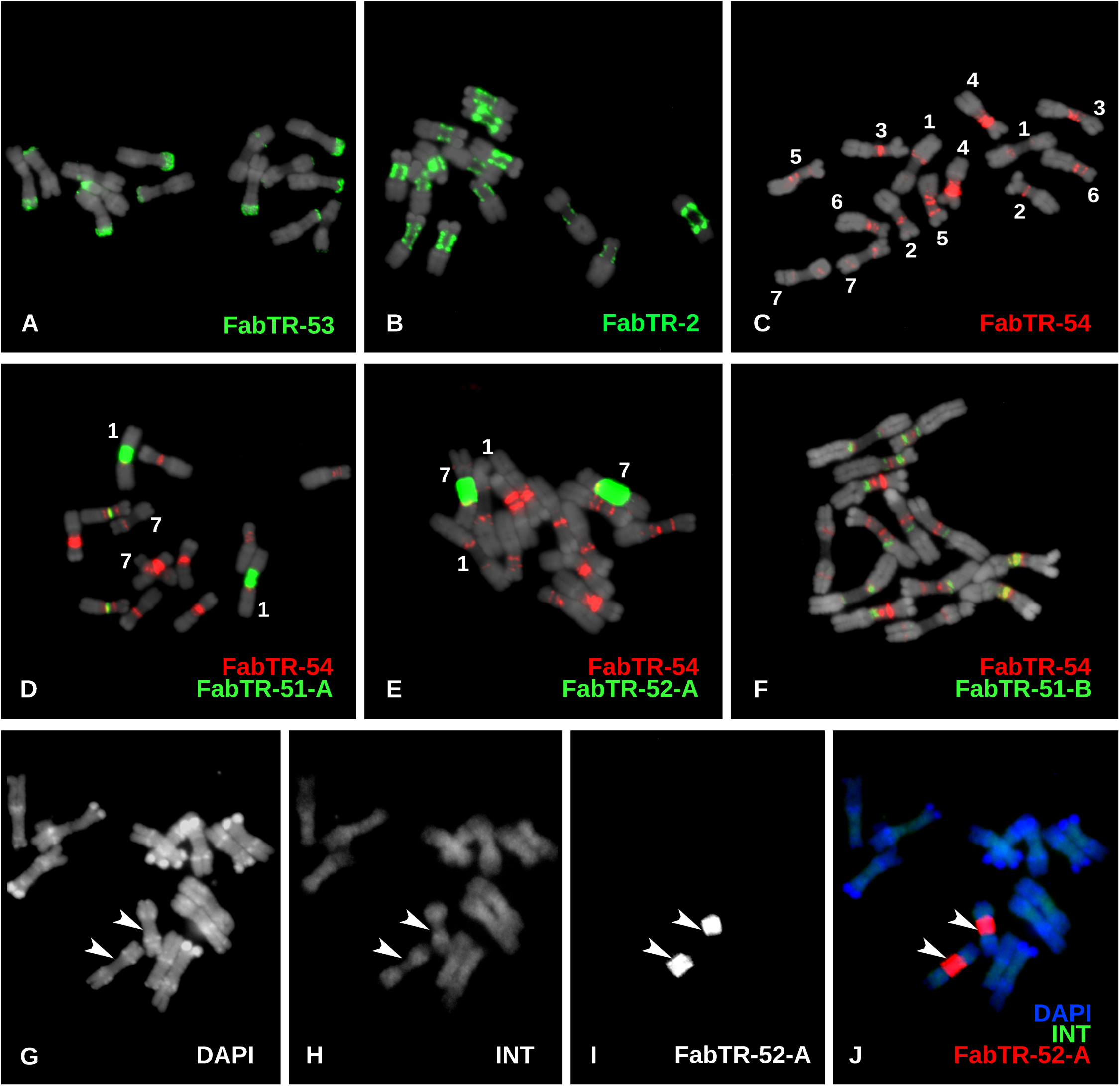
Distribution of the satellite repeats on the metaphase chromosomes of *L. sativus* (2n = 14). (**A-F**) The satellites were visualized using multi-color FISH, with individual probes labeled as indicated by the color-coded descriptions. The chromosomes counterstained with DAPI are shown in gray. The numbers in panel (**C**) correspond to the individual chromosomes that were distinguished using the hybridization patterns of the FabTR-54 sequences. This satellite was then used for chromosome discrimination in combination with other probes. (**G-I**) Simultaneous detection of the Ogre integrase probe (INT) and the satellite FabTR-52 subfamily A demonstrates the different distribution of these sequences in the genome. The probe signals and DAPI counterstaining are shown as separate grayscale images (**G-I**) and a merged image (**J**). The arrows point to the primary constrictions of chromosomes 7.

Although the FISH signals of the Ogre-derived satellites were supposed to originate from their expanded and sequence-homogenized arrays, we had to consider the possibility that the probes had also cross-hybridized to the short repeat arrays within the elements; therefore these FISH patterns may have reflected the genome distribution of Ogre elements. Thus, we investigated the Ogre distribution in the *L. sativus* genome using a probe designed from the major sequence variant of the integarse coding domain of the elements carrying the satellite repeats (see the element scheme in Fig. 4B). The probe produced signals dispersed along the whole chromosomes that differed from the locations of the bands in the primary constrictions revealed by the satellite repeat probes (Fig. 6G-I). Thus, these results confirmed that, while the Ogre elements carrying short tandem repeat arrays were dispersed throughout the genome, these arrays expanded and gave rise to long satellite arrays only within the primary constrictions.

## Discussion

In this work, we demonstrated that the detection and analysis of satellite repeat arrays in the bulk of individual nanopore reads is an efficient method to characterize satellite DNA properties in a genome-wide manner. This is a new addition to an emerging toolbox of approaches utilizing long sequence reads for investigating satellite DNA in complex eukaryotic genomes. Currently, these approaches have primarily been based on generating improved assemblies of satellite-rich regions and their subsequent analyses (Weissensteiner *et al*., 2017; Jain *et al*., 2018). Alternatively, satellite array length variation was analyzed using the long reads aligned to the reference genome (Mitsuhashi *et al*., 2019) or by detecting a single specific satellite locus in the reads (Roeck *et al*., 2018). Compared to these approaches, our strategy does not distinguish individual satDNA arrays in the genome. Instead, our approach applies statistics to partial information gathered from individual reads to infer the general properties of the investigated repeats. As such, this approach can analyze any number of different satellite repeats simultaneously and without the need for a reference genome. However, the inability to specifically address individual repeat loci in the genome may be considered a limitation of our approach. For example, we could not precisely measure the sizes of the arrays that were longer than the analyzed reads and instead provided lower bounds of their lengths. On the other hand, we could reliably distinguish tandem repeats that occurred in the genome predominantly in the form of short arrays from those forming only long contiguous arrays and various intermediate states between these extremes. Additionally, we could analyze the internal arrangements of the identified arrays and characterized the sequences that frequently surrounded the arrays in the genome. This analysis was achieved with a sequencing coverage that was substantially lower compared with that needed for genome assembly. Thus, this approach could be of particular use when analyzing very large genomes, genomes of multiple species in parallel or simply whenever sequencing resources are limited.

We found that only two of the eleven-most abundant satellite repeats occurred in the genome exclusively as long tandem arrays typical of satellite DNA. Both occupied specific genome regions, FabTR-2 was associated with centromeric chromatin, and FabTR-53 made up subtelomeric heterochromatic bands on mitotic chromosomes. Both are also present in other *Fabeae* species (Macas *et al*., 2015), which suggests that they are phylogenetically older compared with the rest of the investigated *L. sativus* satellites. The other feature common to these satellites was the occurrence of HORs that emerge when a satellite array becomes homogenized by units longer than single monomers. The factors that trigger this shift are not clear, however, it is likely that chromatin structure plays a role in this process by exposing only specific, regularly-spaced parts of the array to the recombination-based homogenization. There are examples of HORs associated with specific types of chromatin (Henikoff *et al*., 2015) or chromosomal locations (Macas *et al*., 2006), but data from a wider range of species and diverse satellite repeats are needed to provide a better insight into this phenomenon. The methodology presented here may be instrumental in this task because both the fast Fourier transform and the nucleotide autocorrelation function algorithms employed for the periodicity analyses proved to be accurate and capable of processing large volume of sequence data provided by nanopore sequencing.

One of the key findings of this study is that the majority of *L. sativus* satellites originated from short tandem repeats present in the 3’ untranslated regions (3’UTRs) of Ogre retrotransposons. These hypervariable regions made of tandem repeats that vary in sequences and lengths of their monomers are common in elements of the Tat lineage of plant LTR-retrotransposons, including Ogres (Macas *et al*., 2009; Neumann *et al*., 2019). These tandem repeats were hypothesized to be generated during element replication by illegitimate recombination or abnormal strand transfers between two element copies that are co-packaged in a single virus-like particle (Macas *et al*., 2009); however, the exact mechanism is yet to be determined. The same authors also documented several cases of satellite repeats that likely originated by the amplification of 3’UTR tandem repeats. In addition to proving this mechanism by detecting various stages of the retroelement array expansions in the nanopore reads, the present work on *L. sativus* is the first in which this phenomenon was found to be responsible for the emergence of so many different satellites within a single species. Considering the widespread occurrence and high copy numbers of Tat/Ogre elements in many plant taxa (Neumann *et al*., 2006; Macas & Neumann, 2007; Kubát *et al*., 2014; Macas *et al*., 2015), it can be expected that they play a significant role in satDNA evolution by providing a template for novel satellites that emerge by the expansion of their short tandem repeats. Additionally, similar tandem repeats occur in other types of mobile elements; thus, this phenomenon is possibly even more common. For example, tandem repeats within the DNA transposon *Tetris* have been reported to give rise to a novel satellite repeat in *Drosophila virilis* (Dias *et al*., 2014).

The other important observation presented here is that the long arrays of all nine Ogre-derived satellites are predominantly located in the primary constrictions of metaphase chromosomes. This implies that these regions are favorable for array expansion, perhaps due to specific features of the associated chromatin. Indeed, it has been shown that extended primary constrictions of *L. sativus* carry a distinct type of chromatin that differs from the chromosome arms by the histone phosphorylation and methylation patterns (Neumann *et al*., 2016). However, it is not clear how these chromatin features could promote the amplification of satellite DNA. An alternative explanation could be that the expansion of the Ogre-derived tandem arrays occurs randomly at different genomic loci, but the expanded arrays persist better in the constrictions compared with the chromosome arms. Because excision and eventual elimination of tandem repeats from chromosomes is facilitated by their homologous recombination (Navrátilová *et al*., 2008), this explanation would be supported by the absence of meiotic recombination in the centromeric regions. The regions with suppressed recombination have also been predicted as favorable for satDNA accumulation by computer models (Stephan, 1986). These hypotheses can be tested in the future investigations of properly selected species. For example, the species known to carry chromosome regions with suppressed meiotic recombination located apart from the centromeres would be of particular interest. Such regions occur, for instance, on sex chromosomes (Vyskot & Hobza, 2015), which should allow for assessments of the effects of suppressed recombination without the eventual interference of the centromeric chromatin. In this respect, the spreading of short tandem arrays throughout the genome by mobile elements represents a sort of natural experiment, providing template sequences for satDNA amplification, which in turn, could be used to identify genome and chromatin properties favoring satDNA emergence and persistence in the genome.

## Methods

### DNA isolation and nanopore sequencing

Seeds of *Lathyrus sativus* were purchased from Fratelli Ingegnoli S.p.A. (Milano, Italy, cat.no. 455). High molecular weight (HMW) DNA was extracted from leaf nuclei isolated using a protocol adapted from (Vershinin & Heslop-Harrison, 1998) and (Macas *et al*., 2007). Five grams of young leaves were frozen in liquid nitrogen, ground to a fine powder and incubated for 5 min in 35 ml of ice-cold H buffer (1x HB, 0.5 M sucrose, 1 mM phenylmethylsulphonylfluoride (PMSF), 0.5% (v/v) Triton X-100, 0.1% (v/v) 2-mercaptoethanol). The H buffer was prepared fresh from 10x HB stock (0.1 M TRIS-HCl pH 9.4, 0.8 M KCl, 0.1 M EDTA, 40 mM spermidine, 10 mM spermine). The homogenate was filtered through 48 μmm nylon mesh, adjusted to 35 ml volume with 1x H buffer, and centrifuged at 200 × g for 15 min at 4°C. The pelleted nuclei were resuspended and centrifuged using the same conditions after placement in 35 ml of H buffer and 15 ml of TC buffer (50 mM TRIS-HCl pH 7.5, 75 mM NaCl, 6 mM MgCl_2_, 0.1 mM CaCl_2_). The final centrifugation was performed for 5 min only, and the nuclei were resuspended in 2 ml of TC. HMW DNA was extracted from the pelleted nuclei using a modified CTAB protocol (Murray & Thompson, 1980). The suspension of the nuclei was mixed with an equal volume of 2x CTAB buffer (1.4 M NaCl, 100 mM Tris-HCl pH 8.0, 2% CTAB, 20 mM EDTA, 0.5% (w/v) Na_2_S_2_O_5_, 2% (v/v) 2-mercaptoethanol) and incubated at 50°C for 30-40 min. The solution was extracted with chloroform : isoamylalcohol (24:1) using MaXtract^TM^ High Density Tubes (Qiagen) and precipitated with a 0.7 volume of isopropanol using a sterile glass rod to collect the DNA. Following two washes in 70% ethanol, the DNA was dissolved in TE and treated with 2 μml of RNase Cocktail^TM^ Enzyme Mix (Thermo Fisher Scientific) for 1 h at 37°C. The DNA integrity was checked by running a 200 ng aliquot on inverted field gel electrophoresis (FIGE Mapper, BioRad). Because intact HMW DNA gave poor yields when used with the Oxford Nanopore Ligation Sequencing Kit, the DNA was mildly fragmented by slowly passing the sample through a 0.3 × 12 mm syringe to get a fragment size distribution ranging from ∼30 kb to over 100 kb. Finally, the DNA was further purified by mixing the sample with a 0.5 volume of CU and a 0.5 volume of IR solution from the Qiagen DNeasy PowerClean Pro Clean Up Kit (Qiagen), centrifugation for 2 min at 15,000 rpm at room temperature and DNA precipitation from the supernatant using a 2.5 volume of 96% ethanol. The DNA was dissolved in 10 mM TRIS-HCl pH 8.5 and stored at 4°C.

The sequencing libraries were prepared from 3 μg of the partially fragmented and purified DNA using a Ligation Sequencing Kit SQK-LSK109 (Oxford Nanopore Technologies) following the manufacturer’s protocol. Briefly, the DNA was treated with 2 μml of NEBNext FFPE DNA Repair Mix and 2 μml of NEBNext Ultra II End-prep enzyme mix in a 60 μml volume that also included μml of FFPE and 3.5 μml of End-prep reaction buffers (New England Biolabs). The reaction was performed at 20°C for 5 min and 65°C for 5 min. Then, the DNA was purified using a 0.4x volume of AMPure XP beads (Beckman Coulter); because long DNA fragments caused clumping of the beads and were difficult to detach, the elution was performed with 3 mM TRIS-HCl (pH 8.5) and was extended up to 40 min. Subsequent steps including adapter ligation using NEBNext Quick T4 DNA Ligase and the library preparation for the sequencing were performed as recommended. The whole library was loaded onto FLO-MIN106 R9.4 flow cell and sequenced until the number of active pores dropped below 40 (21-24 h). Two sequencing runs were performed, and the acquired sequence data was first analyzed separately to examine eventual variations. However, because the runs generated similar read length profiles and analysis results, the data were combined for the final analysis.

### Bioinformatic analysis of the nanopore reads

The raw nanopore reads were basecalled using Oxford Nanopore basecaller Guppy (ver. 2.3.1). Quality-filtering of the resulting FastQ reads and their conversion to the FASTA format were performed with BBDuk (part of the BBTools, https://jgi.doe.gov/data-and-tools/bbtools/) run with the parameter maq=8. Reads shorter than 30 kb were discarded. Unless stated otherwise, all bioinformatic analyses were implemented using custom Python and R scripts and executed on a Linux-based server equipped with 64 GB RAM and 32 CPUs.

Satellite repeat sequences were detected in the nanopore reads by similarity searches against a reference database compiled from contigs assembled from clusters of *L. sativus* Illumina reads in the frame of our previous study (Macas *et al*., 2015). Additionally, the database included consensus sequences and their most abundant sequence variants calculated from the same Illumina reads using the TAREAN pipeline (Novák *et al*., 2017) executed with the default parameters and cluster merging option enabled. For each satellite, the reference sequences in the database were placed in the same orientation to allow for the evaluation of the orientations of the satellite arrays in the nanopore reads. The sequence similarities between the reads and the reference database were detected using LASTZ (Harris, 2007). The program parameters were fine-tuned for error-prone nanopore reads using a set of simulated and real reads with known repeat contents while employing visual evaluation of the reported hits using the Integrative Genomics Viewer (Thorvaldsdóttir *et al*., 2013). The LASTZ command including the optimized parameters was “lastz nanopore_reads[multiple, unmask] reference_database-format=general: name1,size1,start1,length1,strand1,name2,size2,start2,length2,strand2,identity, score –ambiguous =iupac --xdrop=10 --hspthresh=1000”. Additionally, the hits with bit scores below 7000 and those with lengths exceeding 1.23x the length of the corresponding reference sequence were discarded (the latter restriction was used to discard the partially unspecific hits that spanned a region of unrelated sequence embedded between two regions with similarities to the reference). Because the similarity searches typically produced large numbers of overlapping hits, they were further processed using custom scripts to detect the coordinates of contiguous repeat regions in the reads (Fig. 1). The regions longer than 300 bp (satellite repeats) or 500 bp (rDNA and telomeric repeats) were recorded and further analyzed. The positions and orientations of the detected satellites were recorded in the form of coded reads where nucleotide sequences were replaced by characters representing the codes for the detected repeats and their orientations, or “0” and “X”, which denoted no detected repeats and annotation conflicts, respectively. In the case of the analysis of repeats other than satellites, the reference databases were augmented for assembled contig sequences representing the following most abundant groups of *L. sativus* dispersed repeats: Ty3/gypsy/Ogre, Ty3/gypsy/Athila, Ty3/gypsy/Chromovirus, Ty3/gypsy/other, Ty1/copia/Maximus, Ty1/copia/other, LTR/unclassified and DNA transposon. These repeats were not arranged nor scored with respect to their orientations. In cases of annotation conflicts of these repeats with the selected satellites, they were scored with lower priority.

Detection of the retrotransposon protein coding domains in the read sequences was performed using DANTE, which is a bioinformatic tool available on the RepeatExplorer server (https://repeatexplorer-elixir.cerit-sc.cz/) employing the LAST program (Kielbasa *et al*., 2011) for similarity searches against the REXdb protein database (Neumann *et al*., 2019). The hits were filtered to pass the following cutoff parameters: minimum identity = 0.3, min. similarity = 0.4, min. alignment length = 0.7, max. interruptions (frameshifts or stop codons) = 10, max. length proportion = 1.2, and protein domain type = ALL. The positions of the filtered hits were then recorded in coded reads as described above.

Analysis of the association of the satellite arrays with other repeats was performed by summarizing the frequencies of all types of repeats detected within 10 kb regions directly adjacent to all arrays of the same satellite repeat family. Visual inspection of the repeat arrangement within the individual nanopore reads using self-similarity dot-plot analysis was performed using the Dotter (Sonnhammer & Durbin, 1995) and Gepard (Krumsiek *et al*., 2007) programs.

Periodicity analysis was performed for the individual satellite repeat arrays longer than 30 kb that were extracted from the nanopore reads and plotted for each array separately or averaged for all arrays of the same satellite. The analysis was performed using the fast Fourier transform algorithm (Venables & Ripley, 2002) as implemented in R programming environment. Briefly, a nucleotide sequence *X* was converted to its numerical representation 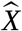 where

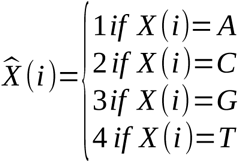

For the resulting sequences of integers, fast Fourier transform was conducted, and the frequencies *f* from the frequency spectra were converted to periodicity *T* as:

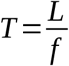

where *L* is the length of the analyzed satellite array. The analysis reveals the lengths of monomers and other tandemly repeated units like HORs as peaks at the corresponding positions on the resulting periodicity spectrum. However, it should be noted that, while these sequence periodicities will always be represented by peaks, some additional peaks with shorter periods could have merely reflected higher harmonics that are present due to the non-sine character of the numerical representation of nucleotide sequences (Li, 1997; Sharma *et al*., 2004). Alternatively, periodicity was analyzed using the autocorrelation function as implemented in the R programming environment (McMurry & Politis, 2010). Nucleotide sequence, X, was first converted to four numerical representations: 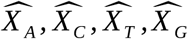 where:

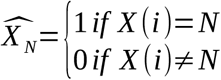

The resulting numerical series were used to calculate the autocorrelations with a lag ranging from 2 to 2000 nucleotides.

### Chromosome preparation and fluorescence in situ hybridization (FISH)

Mitotic chromosomes were prepared from root tip meristems synchronized using 1.18 mM hydroxyurea and 15 μM oryzalin as described previously (Neumann *et al*., 2015). Synchronized root tip meristems were fixed in a 3:1 v/v solution of methanol and glacial acetic acid for 2 days at 4°C. Then the meristems were washed in ice-cold water and digested in 4% cellulase (Onozuka R10, Serva Electrophoresis, Heidelberg, Germany), 2% pectinase and 0.4% pectolyase Y23 (both MP Biomedicals, Santa Ana, CA) in 0.01 M citrate buffer (pH 4.5) for 90 min at 37°C. Following the digestion, the meristems were carefully washed in ice-cold water and postfixed in the 3:1 fixative solution for 1 day at 4°C. The chromosome spreads were prepared by transferring one meristem to a glass slide, macerating it in a drop of freshly made 3:1 fixative and placing the glass slide over a flame as described in (Dong *et al*., 2000). After air-drying, the chromosome preparation were kept at −20°C until used for FISH.

Oligonucleotide FISH probes were labeled with biotin, digoxigenin or rhodamine-red-X at their 5’ ends during synthesis (Integrated DNA Technologies, Leuven, Belgium). They were used for all satellite repeats except for FabTR-53, for which two genomic clones, c1644 and c1645, were used instead. The clones were prepared by PCR amplification of *L. sativus* genomic DNA using primers LASm7c476F (5’-GTT TCT TCG TCA GTA AGC CAC AG-3’) and LASm7c476R (5’-TGG TGA TGG AGA AGA AAC ATAT TG-3’), cloning the amplified band and sequence verification of randomly picked clones as described (Macas *et al*., 2015). The same approach was used to generate probe corresponding to the integrase coding domain of the Ty3/gypsy Ogre elements. The PCR primers used to amplify the prevailing variant A (clone c1825) were PN_ID914 (5’-TCT CMY TRG TGT ACG GTA TGG AAG-3’) and PN_ID915 (5’-CCT TCR TAR TTG GGA GTC CA-3’). The sequences of all probes are provided in Supplementary file 2. The clones were biotin-labeled using nick translation (Kato *et al*., 2006). FISH was performed according to (Macas *et al*., 2007) with hybridization and washing temperatures adjusted to account for the AT/GC content and hybridization stringency while allowing for 10-20% mismatches. The slides were counterstained with 4′,6-diamidino-2-phenylindole (DAPI), mounted in Vectashield mounting medium (Vector Laboratories, Burlingame, CA) and examined using a Zeiss AxioImager.Z2 microscope with an Axiocam 506 mono camera. The images were captured and processed using ZEN pro 2012 software (Carl Zeiss GmbH).

## Availability of source code and requirements

- Project Name: nanopore-read-annotation
- Project homepage: https://github.com/vondrakt/nanopore-read-annotation
- Operating system(s): Linux
- Programming language: python3, R
- Other requirements: R packages: TSclust, Rfast, Biostrings (Bioconductor),
- License: GPLv3

## Availability of supporting data and materials

Raw nanopore reads are available in the European Nucleotide Archive (https://www.ebi.ac.uk/ena) under run accession numbers ERR3374012 and ERR3374013.

## Declarations

## List of abbreviations

aRH: archeal ribonuclease H
FISH: fluorescence *in situ* hybridization
HMW: high molecular weight
HOR: higher order repeat
INT: integrase
LTR: long terminal repeat
PROT: protease
RH: ribonuclease H
RT: reverse transcriptase
satDNA: satellite DNA.

## Consent for publication

Not applicable.

## Competing interests

The authors declare that they have no competing interests.

## Funding

This work was supported supported by the ERDF/ESF project ELIXIR-CZ - Capacity building (No. CZ.02.1.01/0.0/0.0/16_013/0001777) and the ELIXIR-CZ research infrastructure project (MEYS No: LM2015047) including access to computing and storage facilities.

## Authors’ contributions

J.M. conceived the study and drafted the manuscript. T.V. and P.No. developed the scripts for the bioinformatic analysis, and T.V., P.No., P.Ne. and J.M. analyzed the data. A.K. isolated the HMW genomic DNA and cloned the FISH probes. J.M. performed the nanopore sequencing.

L.A.R. conducted the FISH experiments. All authors reviewed and approved the final manuscript.

## Acknowledgements

We thank Ms. Vlasta Tetourová and Ms. Jana Látalová for their excellent technical assistance.

---

**Supplementary Fig. S1.**
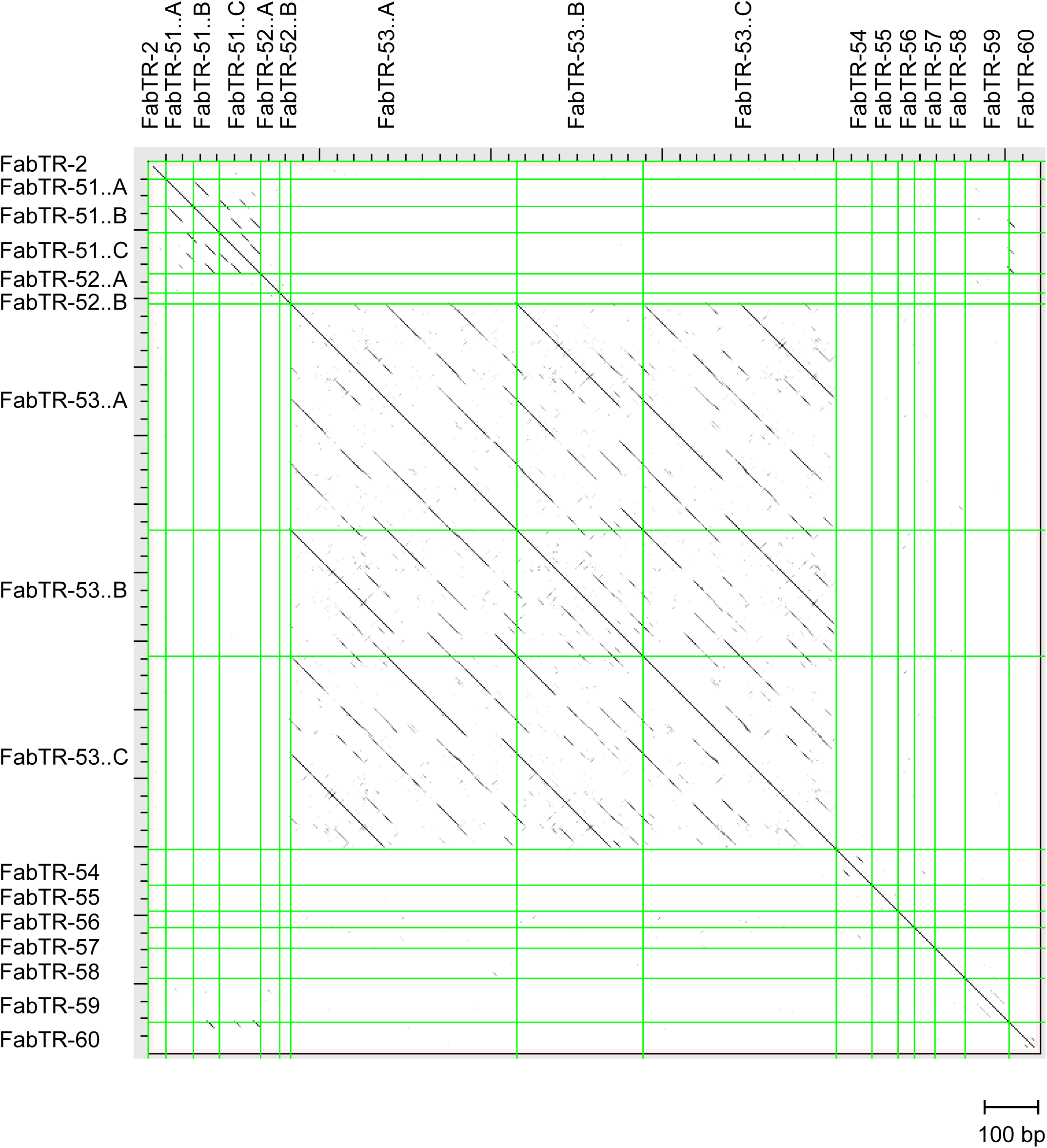
Dot-plot sequence similarity comparison of consensus monomer sequences. The sequences are separated by green lines and their similarities exceeding 40% over a 100 bp sliding window are displayed as black dots or diagonal lines.

**Supplementary Fig. S2.**
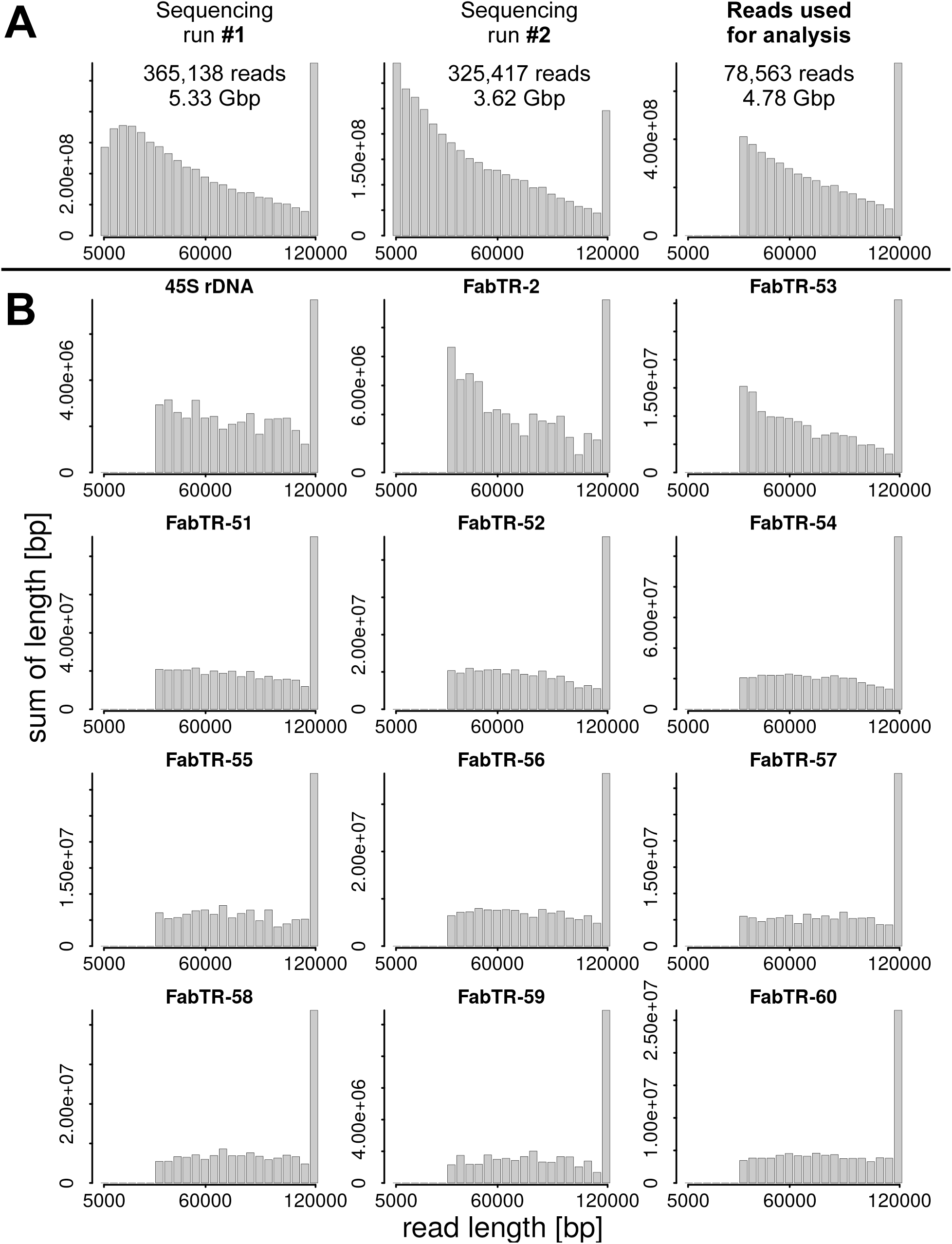
Length distributions of nanopore reads displayed as weighted histograms with bin size of 5 kb, with the last bin including all reads longer than 120 kb. (**A**) Length distributions of raw reads from two sequencing runs and the final set of quality-filtered and size-selected (>30kb) reads used for analysis. (**B**) Length distributions of nanopore reads containing rDNA and satellite repeats.

**Supplementary Fig. S3.**
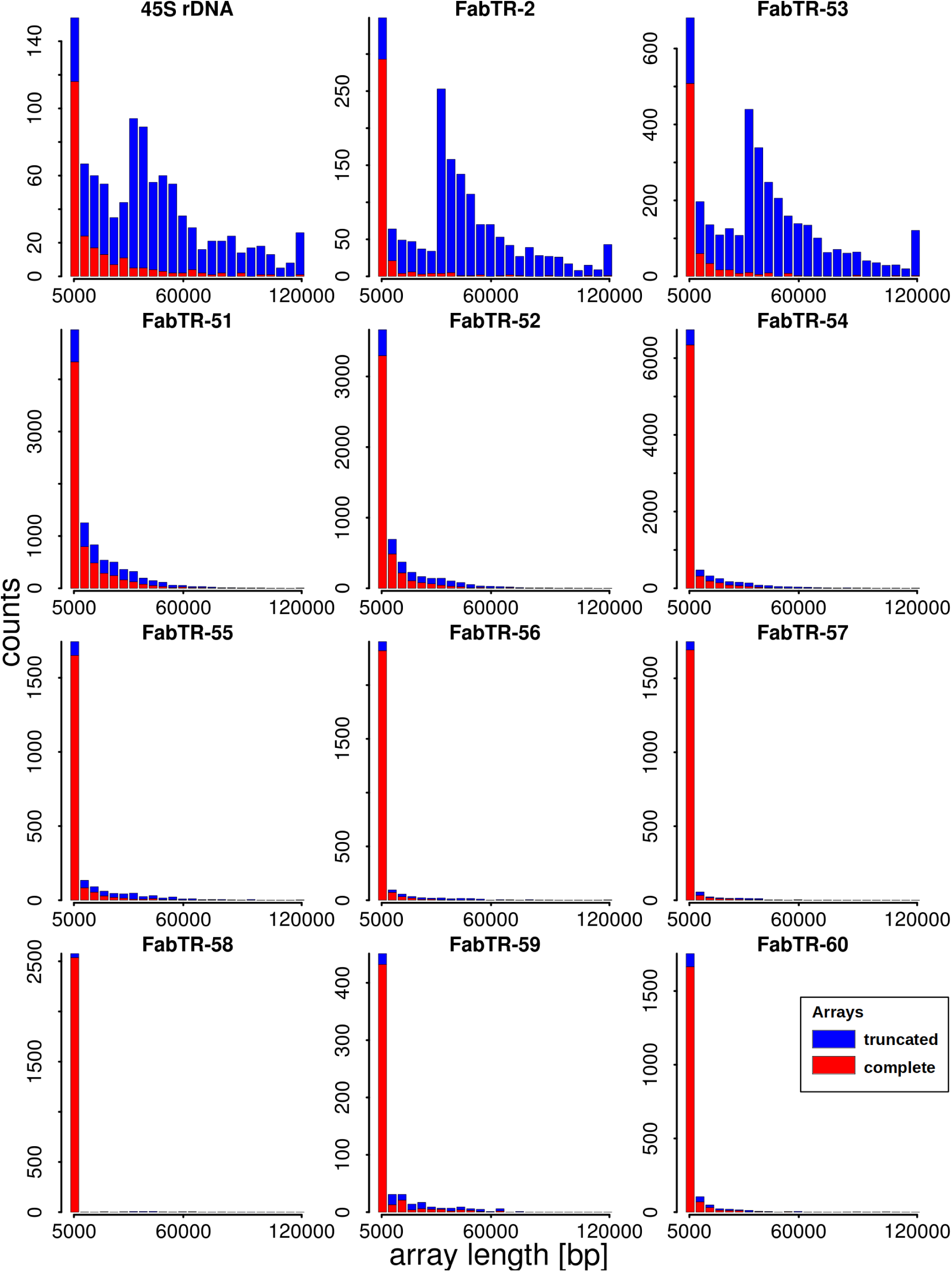
Length distributions of satellite repeat arrays displayed as histograms with bin size of 5 kb, with the last bin including all arrays longer than 120 kb. Arrays which were completely embedded within the reads (red bars) are distinguished from those truncated due to their positions at the ends of the reads (blue bars).

**Supplementary Fig. S4 A-D.**
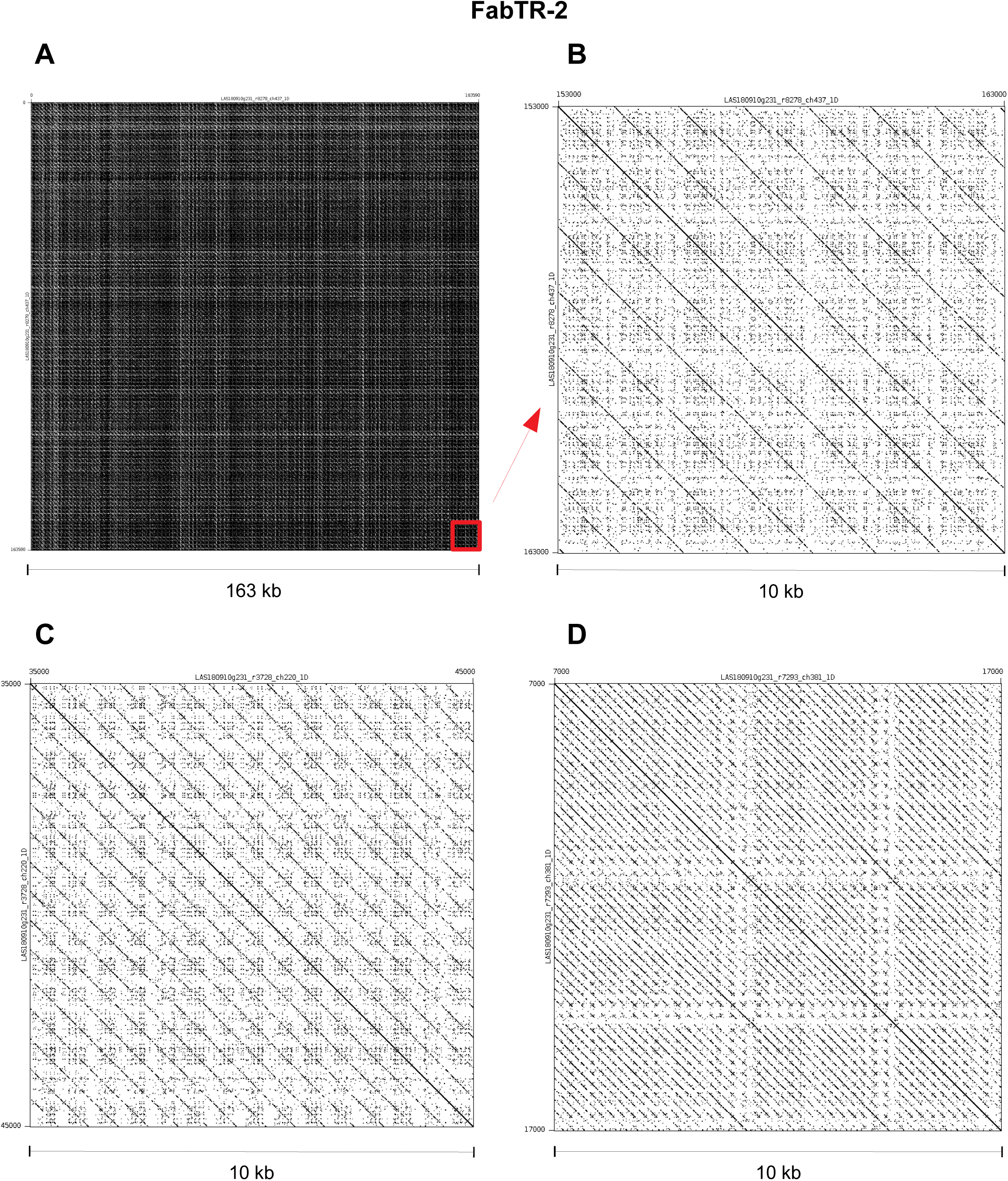
Self-similarity dot-plot visualization of FabTR-2 arrays. Tandem repeats are revealed as diagonal lines with spacing corresponding to monomer length. (**A**) Example of a 163 kb read completely made of FabTR-2 array (the periodicity pattern is obscured by the high density of lines). (**B**) Magnification of the 10 kb region highlighted by a red square on panel A. This array is homogenized as ∼1300 bp HOR. (**C,D**) Examples of other FabTR-2 periodicities detected in different reads (only 10 kb regions were used for dot-plots to make periodicity patterns comparable with other plots).

**Supplementary Fig. S4 E-H.**
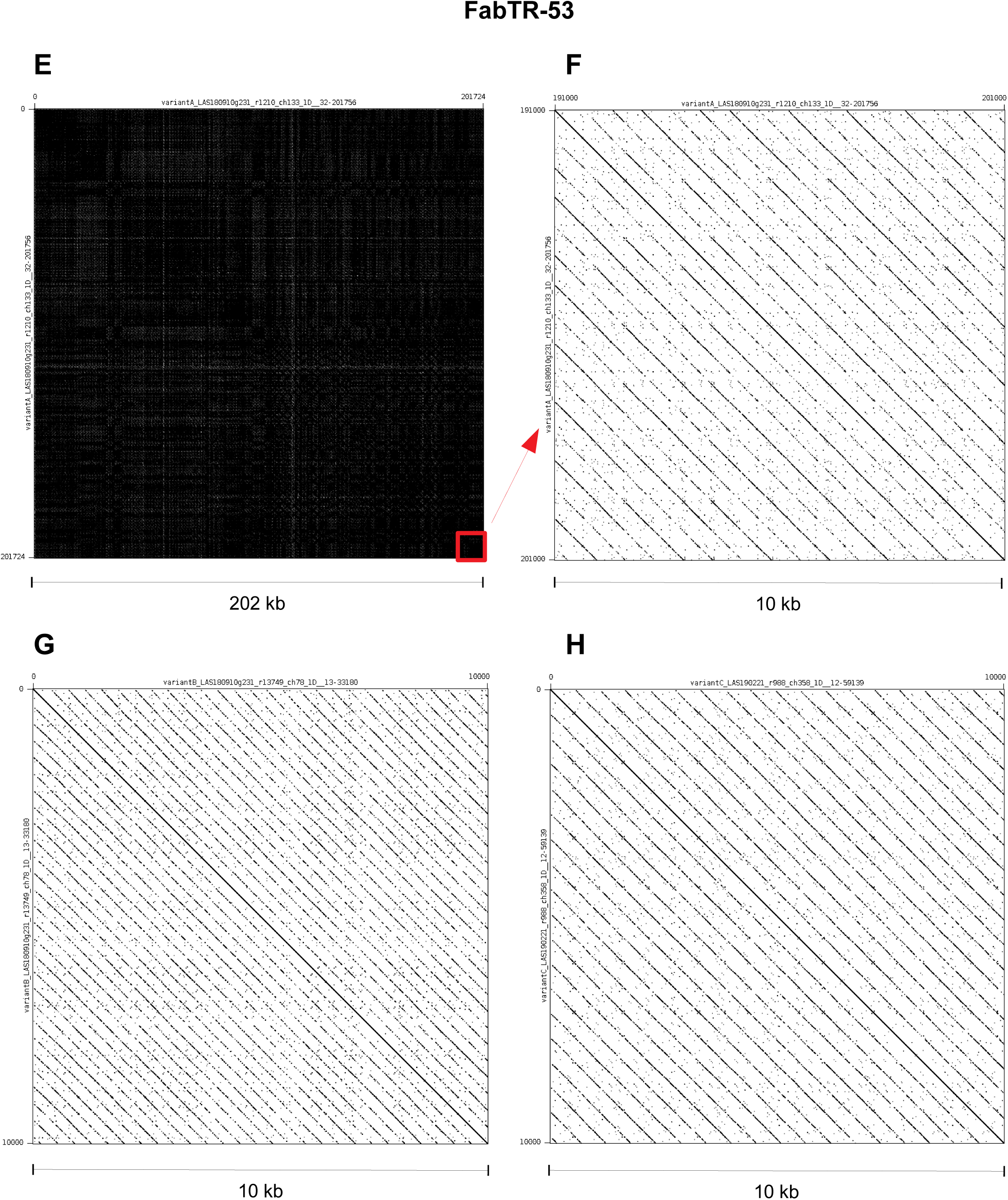
Self-similarity dot-plot visualization of FabTR-53 arrays. (**E**) Example of a 202 kb read completely made of FabTR-2 array (the periodicity pattern is obscured by the high density of lines). (**F**) Magnification of the 10 kb region highlighted by a red square on panel A. (**G,H**) Examples of other FabTR-53 periodicities detected in different reads (only 10 kb regions were used for dot-plots to make periodicity patterns comparable with other plots).

**Supplementary Fig. S4 I-K.**
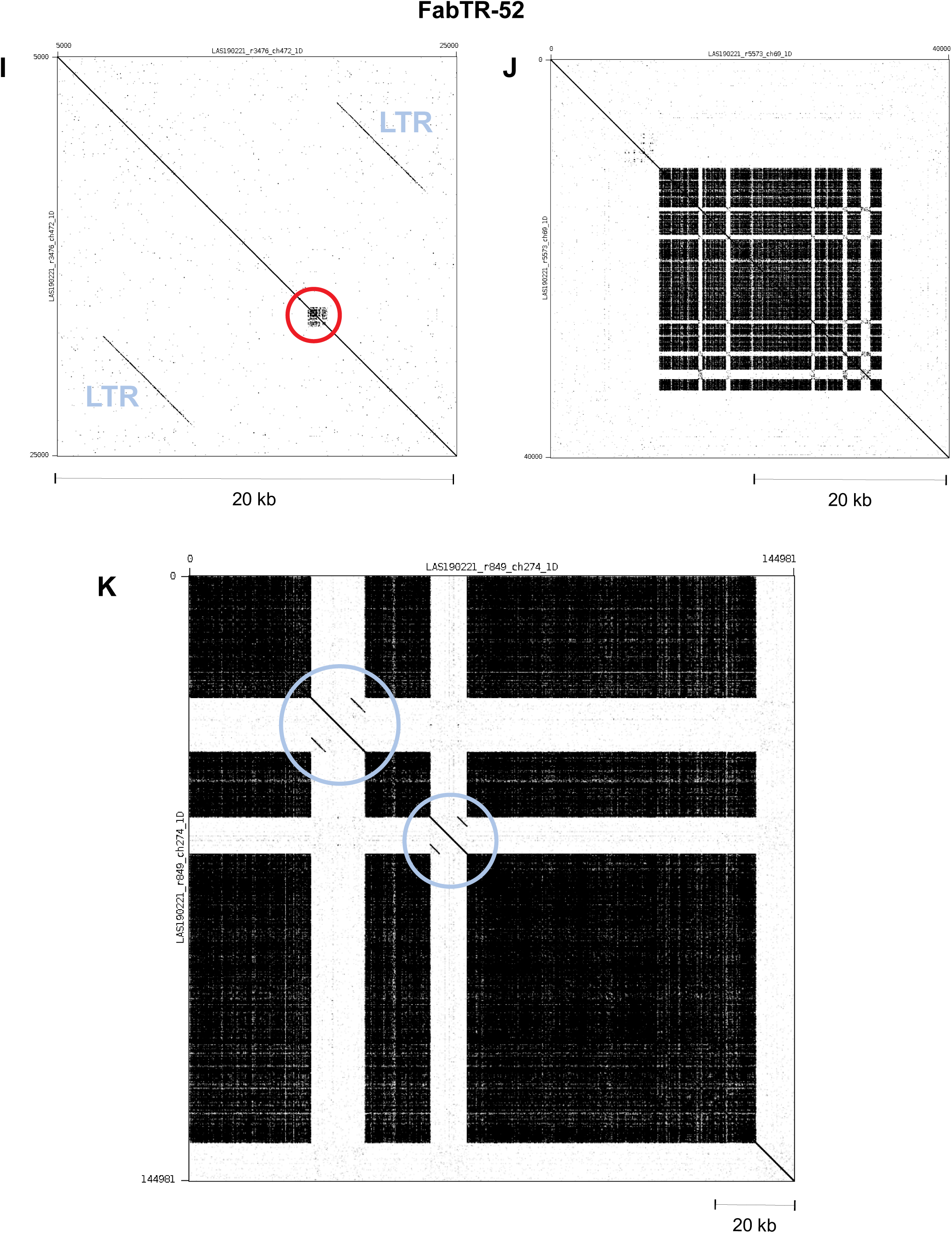
Dot-plots demonstrating length distribution of FabTR-52 arrays, ranging from short arrays (red circle) embedded within LTR-retrotransposon sequences (**I**) and partially expanded arrays (**J**) to the arrays >100 kb in length which are interrupted by insertions of LTR-retrotransposons (blue circles) (**K**).

**Supplementary Fig. S4 L-N.**
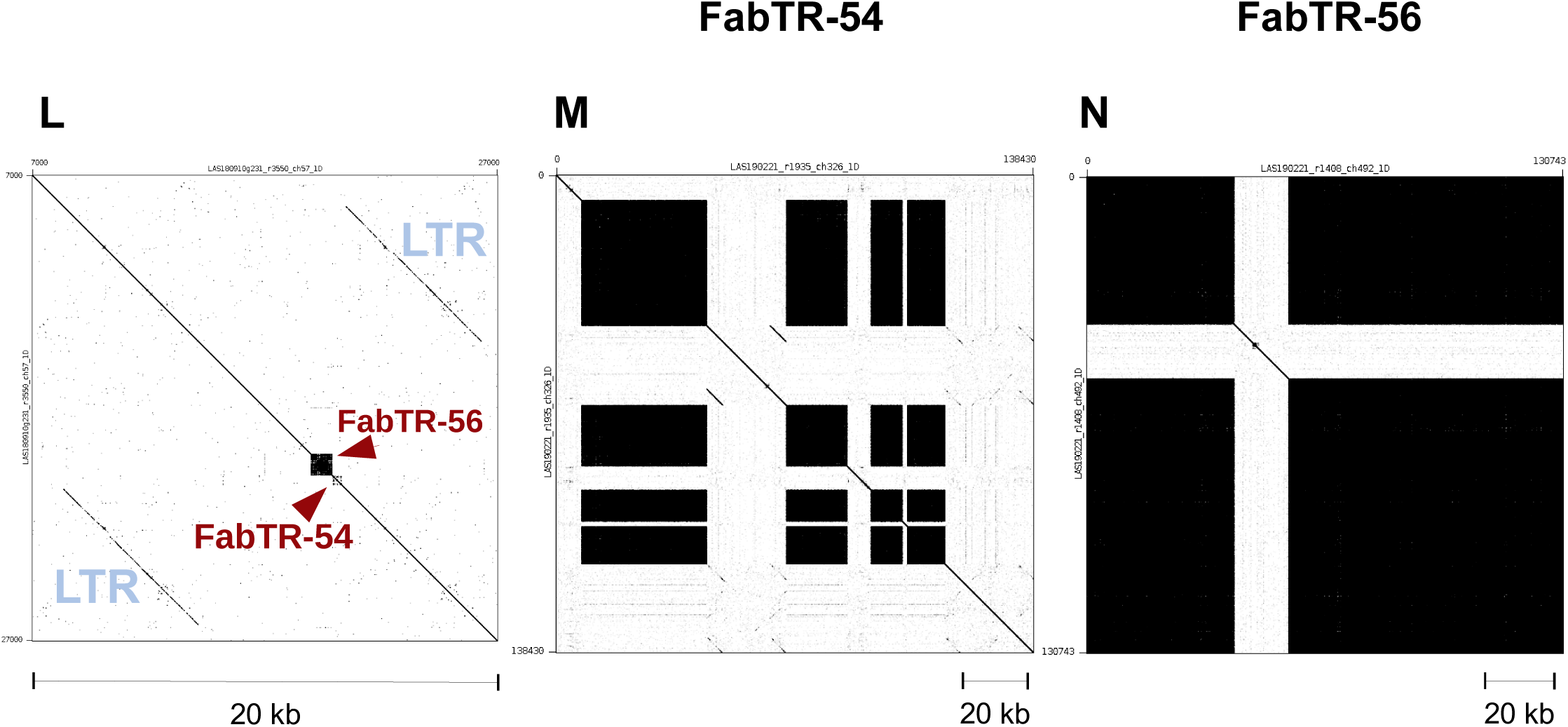
(**L**) Example of LTR-retrotransposon carrying short FabTR-54 and FabTR-56 arrays. Reads with those tandem repeats expanded to long arrays are shown on panels **M** (FabTR-54) and **N** (FabTR-56). The expanded tandem arrays appear as black squares on the dot-plots due to high density of lines.

**Supplementary Fig. S4 O-Q.**
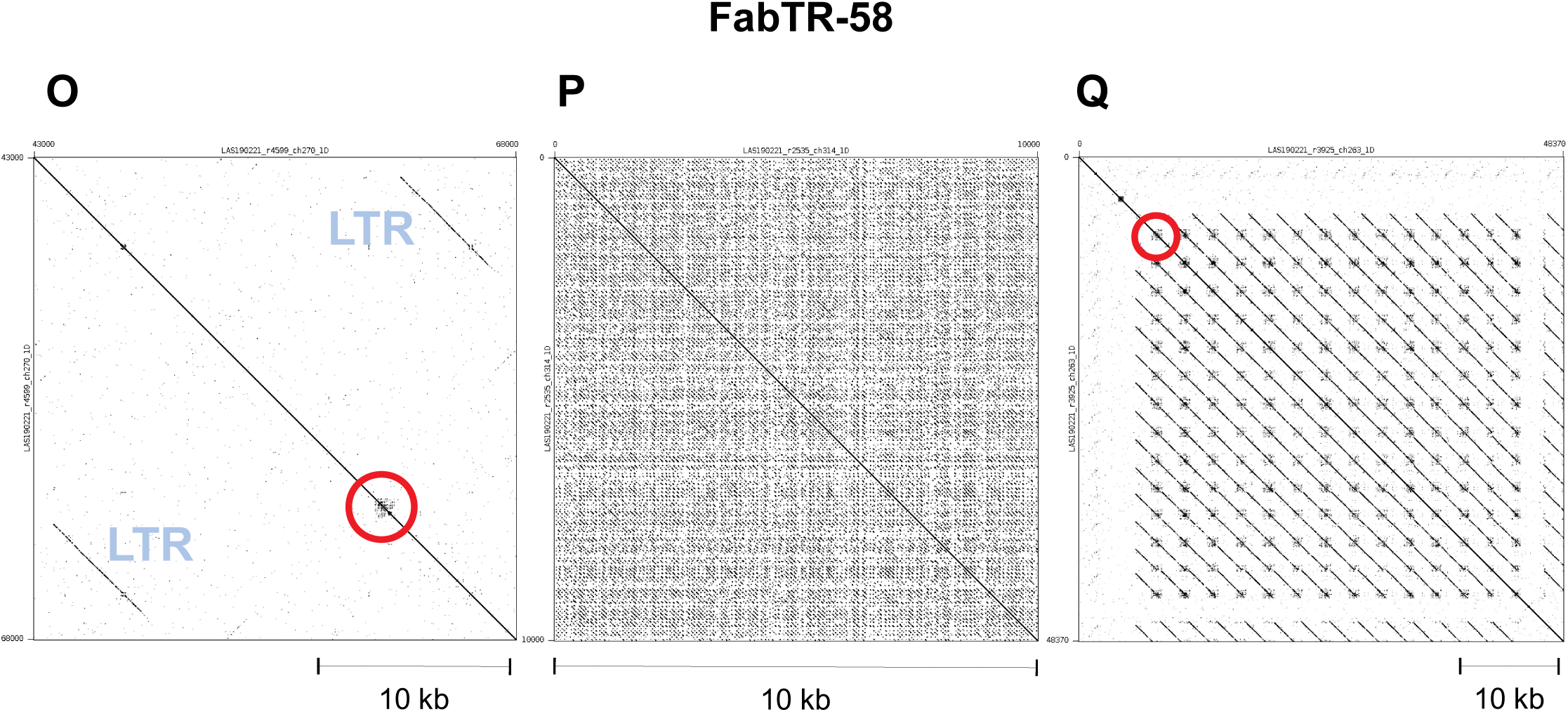
Three types of genome organization of FabTR-58 repeats: (O) short array (marked by red circle) within LTR-retrotransposon, (P) expanded array, (Q) short arrays embedded within a longer tandem repeat monomer.

**Supplementary Fig. S5.**
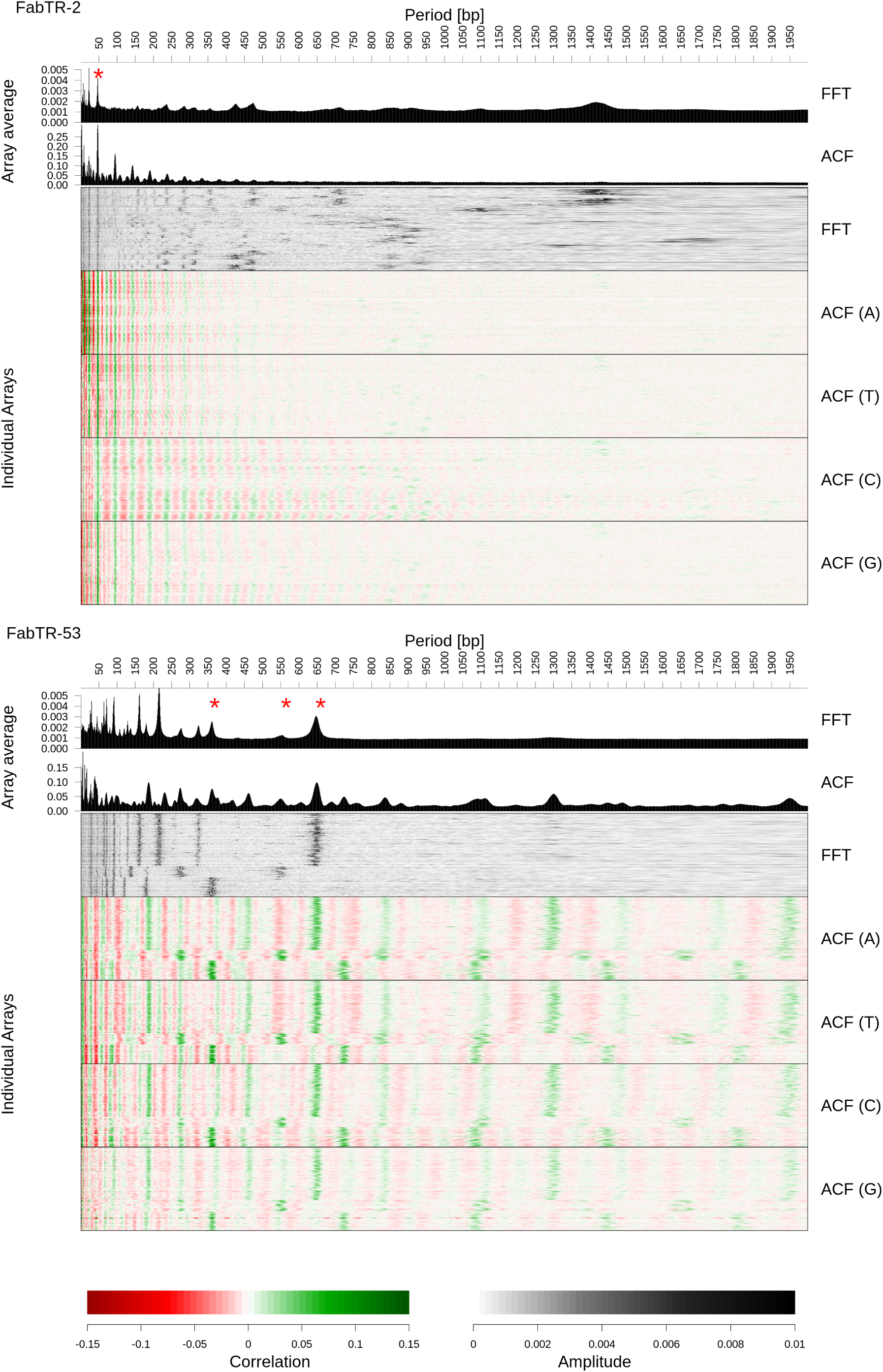
Detailed periodicity analysis of FabTR-2 and FabTR-53 arrays. Periodicity analysis using fast Fourier transform (FFT) and autocorrelation function (ACF) are shown as averages of spectra calculated on individual satellite arrays longer than 30 kb. Periodicity spectra from individual arrays are shown as heatmaps with rows corresponding to individual arrays. Autocorrelations are shown separately for individual nucleotides.

**Supplementary Fig. S6.**
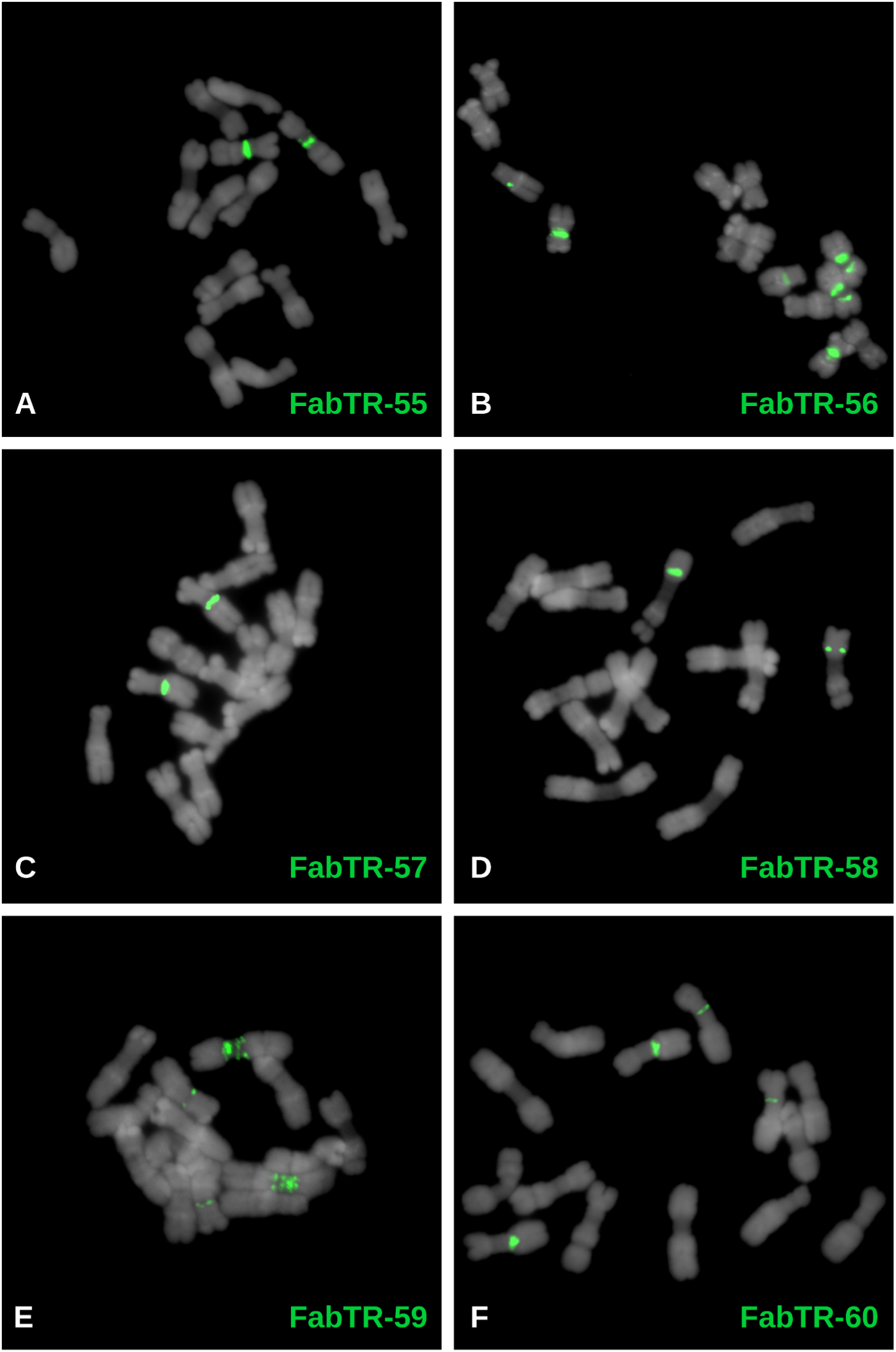
Distribution of the satellite repeats on the metaphase chromosomes of *L. sativus* (2n = 14). The satellites were visualized using FISH, with individual probes labeled as indicated by the color-coded descriptions. The chromosomes counterstained with DAPI are shown in gray.

